# IFNγ-dependent remodelling of the myeloid landscape underlies control of IFNγ-insensitive tumours

**DOI:** 10.1101/2024.03.25.586537

**Authors:** Vivian W.C. Lau, Gracie Mead, Julie M. Mazet, Anagha Krishnan, Edward W. Roberts, Gennaro Prota, Uzi Gileadi, Vincenzo Cerundolo, Audrey Gérard

**Author notes:** Deceased (Vincenzo Cerundolo). Corresponding author: Audrey Gérard.

## Abstract

Loss of IFNγ-sensitivity by tumours is thought to be a mechanism enabling evasion, as some cancers lacking IFNγ-signalling demonstrate resistance to checkpoint immunotherapy. However, recent studies demonstrated that IFNγ-resistant tumours are well-controlled and sensitized for immunotherapy. The underlying mechanism leading to enhanced immune responses in those patients is unknown. Using IFNγ-insensitive melanoma tumours which were well-controlled by the endogenous anti-tumour response, we found that despite low basal MHC class I expression by tumours, CD8^+^ T cell infiltration was not hindered and, unexpectedly, their production of IFNγ was still important for tumour control. Mechanistically, IFNγ triggers pro-inflammatory remodelling of IFNγ-insensitive tumours, affecting the differentiation of myeloid cells. Predominantly, immunosuppressive macrophages are inhibited, while inflammatory phenotypes of monocytes and ‘mono-macs’ are preserved in IFNγ-insensitive tumours. This is supported by a co-dependency between CD8^+^ T cells and monocyte/macrophages, as depletion of one resulted in loss of the other. Our work demonstrates an important mechanistic understanding of how IFNγ resistance does not preclude failure of anti-tumour responses. Importantly, immune remodelling appears to be dominant in IFNγ-sensitive and IFNγ-insensitive mixed tumours, and is enriched in humans with tumours mutated in the IFNγ pathway, suggesting this may be leveraged for therapy in the future.

## Introduction

Tumour escape is a mechanism whereby tumours acquire genetic mutations resulting in the evasion of immunosurveillance. Immune evasion can occur during primary or acquired resistance and is associated with a lack of therapeutic response and subsequent disease progression. Establishment of resistance generally involves loss of T cell-dependent cytotoxicity, which can occur through deficiencies in antigen presentation mechanisms, or acquisition of resistance against interferon gamma (IFNγ)(Kluger et al., 2020; Sharma et al., 2017).

IFNγ induces anti-tumour immunity by exerting direct cytotoxic and cytostatic effects on tumours(Chin et al., 1996; Shankaran et al., 2001), inducing major histocompatibility complex (MHC) expression(Zhou, 2009), and promoting the expansion of effector lymphocytes and maturation of myeloid populations(Pan et al., 2004; Whitmire et al., 2005). But it can simultaneously impede anti-tumour immunity, for example, through induction of PD-L1 and IDO expression(Garcia-Diaz et al., 2017; Spranger et al., 2013) and limiting stem-like T-cell driven immunity(Mazet et al., 2023). As such, IFNγ affects both the tumour itself and its microenvironment, with contrasting consequences on the anti-tumour response(Castro et al., 2018). In patients, it is well-established that durable responses to immune checkpoint blockade (ICB) are associated with IFNγ-related gene signatures(Ayers et al., 2017), which are also correlated with increased tumour mutational burden (TMB)(Fumet et al., 2020) and T cell infiltration(Grasso et al., 2020), suggesting that IFNγ-driven responses is a pre-requisite to response to ICB. IFNγ induces a complex network of downstream effects mediated through its cognate receptor, IFNγR. IFNγ signalling regulates expression of hundreds of genes, known as interferon-stimulated genes (ISGs), and interestingly, the balance between immune and cancer ISGs correlates with response to ICB(Benci et al., 2019), highlighting the importance in eliciting the correct equilibrium between pro- and anti-tumoural IFNγ functions.

Many clinical reports have associated acquired resistance to ICB with loss of IFNγ response by tumour cells via signalling pathway mutations(Gao et al., 2016; Sucker et al., 2017; Zaretsky et al., 2016). Yet, mutations in IFNγ signalling pathway are relatively infrequent (i.e. <10% patients) in colorectal cancers(Shin et al., 2017), and *JAK1* mutations were associated with better 5-year survival rates amongst colorectal cancer patients(Sveen et al., 2017). In patients with frequent mutations such as endometrial cancers, *JAK1* mutations appeared to have little impact on ICB outcomes(Song and Chow, 2023). Recent meta-analyses focused on leveraging statistical power of independent observations have discovered that pre-existing IFNγ pathway mutations in multiple cancer types did not necessitate lack of response to ICB(Chow et al., 2023). Several *in vivo* pre-clinical models and CRISPR screens support this observation, as cell lines with mutations in IFNγR or downstream signalling molecules such as JAK1/2 and STAT1 have been shown to sensitize the tumour towards improved ICB response(Dubrot et al., 2022; Kearney et al., 2018). It is therefore important to understand the factors that contribute to eliciting a potent anti-tumour response during tumour escape and how it integrates with mutations in the IFNγ pathway.

IFNγ signatures pre- and post-ICB treatment are associated with clinical response to therapy, and it was often assumed that the main mode of action of IFNγ is to directly inhibit and/or kill tumour cells. As the above studies challenge this dogma, it is still poorly understood mechanistically what controls tumours insensitive to IFNγ. In this study, we observe substantial remodelling of the tumour immune landscape which favours inflammatory signalling pathways following ablation of IFNγ-signalling on tumour cells. Loss of IFNγR1 on tumour cells in our murine melanoma models results in intra-tumoural IFNγ accumulation, which induces pro-inflammatory signalling of immune-infiltrating populations. The myeloid compartment exhibits substantial immune remodelling in IFNγR1-deficient tumours compared to wild-type, through increased recruitment and retention of pro-inflammatory monocytes and decreased immunosuppressive macrophage generation. More importantly, loss of monocyte infiltration subverts the inflammatory phenotype in IFNγRKO tumours and their intra-tumoural co-localization with CD8^+^ T cells appears to support their anti-tumour functions.

Overall, our study demonstrates that tumour-derived mutations in the IFNγ pathway can trigger a remodelling of the immune landscape underlying the control of those mutated clones. As we also observe this immune landscape remodelling in humans, it highlights the relevance of our findings and provides potential new therapeutic avenues, which could be important beyond IFNγ-insensitive tumours.

## Materials and Methods

### Mice

C57BL/6J (B6) WT male mice from 6 to 8 weeks old were purchased from Charles River (JAX number – 000664) and housed 1-2 weeks before experimentation. IFNγ-GREAT^YFP^ (JAX number – 017580), CD8 KO (JAX number – 002665), IFNγ KO (JAX number – 002287), and CCR2 KO (JAX number – 004999) 6- to 12-week-old male and female mice were housed and bred under specific pathogen-free/SPF conditions in the in-house animal facilities at the University of Oxford. Experimental and control animals were co-housed and kept in individually ventilated cages supplemented with environmental enrichment at 20-24°C, 45-64% humidity, and 12h light/dark cycles. Mice were euthanized by CO_2_ asphyxiation followed by cervical dissociation. All experiments involving mice were conducted in agreement with the United Kingdom Animal Scientific Procedures Act of 1986 and performed in accordance with approved experimental procedures by the Home Office and the Local Ethics Reviews Committee (University of Oxford) under UK project licenses P4BEAEBB5 and PP3609558.

### Cell line generation and culture conditions

B16F10 Tyr^-/-^ expressing mCherry and ovalbumin (B16-OVA) was kindly provided by Dr. Edward Roberts from the Beatson Institute (Glasgow, UK). Knockout of murine IFNGR1 using CRISPR-Cas9-mediated gene editing using protocols described by Ran *et al*. (2013)(Ran et al., 2013). Briefly, the single guide RNA (sgRNA) sequences targeting exon 1 or 2 of murine IFNγR1 were cloned into the pX458 backbone (Addgene) containing Cas9 expression and GFP expression, followed by validation via Sanger sequencing. Target murine cell lines were transiently transfected with pX458 sgIFNγR1 using Lipofectamine 3000 (Invitrogen, Cat. L3000001). After 48 hours, cells were single-cell sorted into individual wells of a 96-well plate. Single cell clones were expanded and stimulated with 1 ng/mL recombinant murine IFNγ (Peprotech, Cat. 315-05). Clones were stained for MHC-I H2-D^b^ expression by flow cytometry analysis. Five individual clones unresponsive to IFNγ stimulation were pooled to form the final cell line used in subsequent experiments. For generation of the cell line expressing IFNγR1 Y445A, wild-type and mutated sequences were synthesized as gBlock gene fragments (Integrated DNA Technologies, Inc.) and cloned into a pLV lentiviral backbone containing puromycin resistance. Wild-type IFNGR1 and Y445A was re-expressed in the B16 IFNγR1KO cell line and selected for puromycin resistant cells. Resulting cell lines were validated by stimulation with rmIFNγ at 10ng/mL for MHC class I re-sensitization. All cell lines were cultured in RPMI 1640 (Gibco, Cat. 21870-076) supplemented with 10% FCS (Sigma, Cat. F9665-500ML), and 1X penicillin/streptomycin/L-glutamine (Gibco, Cat. 10378-016) (referred to as R10 medium). Cell lines were kept at 37°C in 5% CO2 and routinely checked for mycoplasma contamination via LookOut Mycoplasma PCR detection kit (Sigma, Cat. MP0035).

### Tumour induction and administration of immune-modifying agents

Cell lines were harvested at 50-70% confluency on the day of tumour injections using trypsin-EDTA (Sigma, Cat. T3924-500ML) and washed twice in PBS prior to resuspension at desired cell concentrations in PBS or PBS + 25% Matrigel (Corning, Cat. 354262, or Cat. 256231). Mice were anaesthetized using isoflurane (Zoetis) and flanks were shaved prior to injection. Tumours were typically engrafted subcutaneously in the right and/or left ventral flanks at a cell concentration of 1.00-1.25×10^6^ cells per mL, resulting in engraftment of 1×10^5^ or 1.25×10^5^ cells per 100 μL injection. Tumours were measured after 5-7 days post-injection using callipers and monitored for humane endpoints continuously until experimental termination. Tumour volumes were calculated using [(LxWxH)/2] formula in mm^3^.

In some experiments, mice were treated with CD8 or NK1.1 depleting antibodies or isotype controls (BioXCell, anti-CD8β Cat. BE0223, and anti-NK1.1 Cat. BE0036). Mice were injected intraperitonially with 50 µg antibodies every 2 to 3 days and monitored daily.

### Tissue processing

At indicated timepoints, mice were sacrificed and tumours were measured, excised, and weighed before processing. Tumours were dilacerated using scalpels to obtain <1mm sized pieces and resuspended in R10 supplemented with 1 mg/mL Liberase TL (Roche, Cat. 5401020001) and 10 μg/mL DNase I (Roche, Cat. 11284932001) for enzymatic digestion. Tumour suspensions were incubated at 37°C for 30 minutes before physical dissociation of remaining fragments through 70μm cell strainers to obtain single cell suspensions.

### ELISAs and LegendPlex assays

Tumour supernatants were collected before enzymatic digestion following mechanical dilaceration into 1 mL of R10 media. Samples were frozen at −20°C and thawed on ice prior to assaying. IFNγ concentrations were determined using an IFNγ mouse uncoated ELISA kit (Invitrogen, Cat. 88-7314-77) following manufacturer’s protocols. Obtained IFNγ concentrations using the standard curve were normalized to tumour weights. For LEGENDplex™ Mouse Cytokine Release Syndrome Panel (13-plex) assays (Biolegend, Cat. 741024), 25uL of supernatant was used following manufacturer’s protocols and samples were analysed by BD LSRFortessa.

### Flow cytometry

Single cell suspensions from cell lines or tumours were plated into 96-well V bottom plates at concentration of 2.5×10^6^ cells or less. Cells were washed 1X using PBS prior to addition of viability dyes (Biolegend, Zombie NIR, Cat. 423106, or Zombie UV, Cat. 423108) according to manufacturers’ instructions. Samples were incubated with anti-CD16/anti-CD32 blocking antibodies (Biolegend, Cat. 101302) for 20 minutes at 4°C, followed by fluorochrome-conjugated antibodies against extracellular markers for 30 minutes at 4°C. Cells were washed with FACS-EDTA buffer (2% FCS, 2.5 mM EDTA, and 0.01% sodium azide in PBS) and resuspended in 2% paraformaldehyde (Alfa Aesar, Cat. 43368.9M) in PBS for a 20-minute fixation at 4°C. Cells were washed before resuspension in FACS-EDTA buffer and stored until analysis. For experiments where tetramers were used, tetramers were diluted in FACS-EDTA and incubated with samples in-between the Fc-blocking and antibody staining steps. Tetramers were obtained from the NIH Tetramer Core Facility (Atlanta, GA, USA). For samples where calculation of absolute cell numbers was required, 20,000 Polybead polystyrene Black Dyed Microspheres (Polyscience, Cat. 24293-15) were added to the samples prior to analysis. Absolute counts (cells/μL) were calculated as ((Cell count x counting beads volume) / (Counting beads x Cell volume)) x Counting beads concentration (beads/μL). All flow cytometry samples were analysed by BD Fortessa X-20, BD LSRII, or Cytek Aurora as indicated per experiment. Flow cytometry data was analysed using FlowJo V.10 (BD), or OMIQ (Dotmatics).

### Tissue fixation, cryosectioning, and imaging

Whole intact tumours were harvested on days 10-13 post engraftment and immediately immersed in fixative solution (1% PFA, 75 mM L-lysine [Sigma, Cat. L5501], 10 mM sodium m-periodate [Thermo Scientific Pierce, Cat. 20504], diluted in 0.2M PBS adjusted to pH 7.4) for 16-20 hours at 4°C under gentle agitation. Fixative solution was discarded, and tumours were washed using 1X PBS for 1 hour at 4°C under gentle agitation to remove remaining PFA. Tumours were then resuspended in 30% sucrose (w/v, diluted in pH 7.4 PBS) for 24-36 hours at 4°C without agitation, until tissue was no longer floating. Tumours were cryogenically frozen in OCT compound (Thermo Fisher, Cat. 15212776) using methanol and dry ice bath, and stored at - 80°C until cryosectioning. Frozen tumour blocks were cryosectioned at 10 μm thickness using a Leica CM1900UV and mounted onto glass slides (VWR, Cat. 631-0108). Cryosections were stored at −80°C until staining. For staining, sections were washed with PBS to remove OCT compound on the slides, and blocked with solutions containing imaging buffer (2% FCS, 0.1% Triton X-100 (Sigma, Cat. X100), 0.01% sodium azide), FcBlock, and species-specific serum depending on the fluorochrome-conjugated antibodies used in each staining panel. Sections were blocked for a minimum of 3 hours at room temperature, before incubation with fluorescently-conjugated antibodies diluted in blocking solution overnight at 4°C. A final wash was performed twice using imaging buffer before the sections were mounted using Fluoromount G (Southern Biotech, Cat. 0100-01) and glass coverslips were placed on top of the sections. Images were collected using Zeiss Axioscan 7 Slide Scanner or Zeiss LSM 980 confocal microscope, and analysed using Imaris software (Bitplane, V10.0).

### Single-cell RNA sequencing

Single cell suspensions from 3 B16-OVA WT tumours were labelled with TotalSeq(TM)-C0301 antibody (Biolegend, Cat. 155861), and 3 IFNγRKO tumours were labelled with TotalSeq(TM)-C0302 (Biolegend, Cat. 155863). Live cells stained with Zombie NIR and CD45 antibody were sorted based on expression of CD45 using a BD FACSAria™ II. Approximately 10,000 cells per sample were loaded onto the 10X Genomics Chromium Controller (Chip K). Gene expression and feature barcoding libraries were prepared using the 10x Genomics Single Cell 5’ Reagent Kits v2 (Dual Index) following manufacturer user guide (CG000330 Rev B). The final libraries were diluted to ∼10 nM for storage. The 10nM library was denatured and further diluted prior to loading on the NovaSeq6000 sequencing platform (Illumina, v1.5 chemistry, 28 bp/98 bp paired end for gene expression and feature barcoding).

### Analysis of scRNAseq datasets

Sequence reads were mapped using CellRanger multi (version 6.0.0) and the 10x mouse reference transcriptome (version 2020-A). The R package Seurat v4 was used in conjunction with other tools for QC, demultiplexing, filtering, and annotation of the dataset. Briefly, singlets were extracted from the dataset, and counts were log normalized and variable features were scaled. Cells having fewer than 500 or greater than 6000 detected genes were filtered out. Cells in which 5% of the UMIs represent mitochondrial protein-coding genes or more than 20% of large gene contents were also filtered. Lastly, decontX(Yin et al., 2023) was used to determine contamination of droplets with ambient RNA. The filtered dataset was scaled, log normalized, and variable features were identified using the functions in Seurat. Principle component analysis was performed, and the number of PCs used for clustering was determined using the ElbowPlot function. Clusters and markers for clustered was identified using the Louvain algorithm embedded in the FindNeighbours and FindClusters functions, at a resolution of 0.5. UMAP projections were computed using the first 10 principal components. Clusters were annotated using the FindAllMarkers function to determine differentially expressed genes for each cluster, then cluster identities were verified using the package SingleR(Aran et al., 2019). Heatmaps, violin plots, and UMAP projections were generated using Seurat v4(Hao et al., 2021). The FindMarkers function was used to find differentially expressed genes within each cluster between WT and IFNGR1KO conditions. Pathway analysis and plotting of results was performed using the tool fgsea(Korotkevich et al., 2021). Volcano plots and bar plots were created using ggplot2. Module scoring of different TAM subsets was done using the package UCell(Andreatta and Carmona, 2021). Trajectory analysis for CD8^+^ T cell and macrophage clusters was completed and visualized using the package monocle3(Qiu et al., 2017; Trapnell et al., 2014). Finally, analysis of cell-cell communication networks and plotting of results were performed using the package CellChat(Jin et al., 2021).

### Analysis of human datasets

Selected TCGA PanCancer Atlas studies were retrieved from cBioPortal(Cerami et al., 2012; de Bruijn et al., 2023; Gao et al., 2013) and queried for cases which contained gene mutations in IFNγ pathway (IFNGR1, IFNGR2, JAK1, JAK2 and STAT1), antigen presentation pathway (HLA-A, HLA-B, HLA-C, B2M, TAP1 and TAP2), or individual genes as indicated. Survival curves for selected cancer types were also retrieved for patient cases which contained the set of IFNγ-pathway mutations versus cases without such mutations. For enrichment scoring of CD8-monocyte signatures in human cancers, normalized STAR gene counts were retrieved from the TCGA for cases of each cancer type and subdivided into control and mutant groups, where mutants were cases which contained confirmed mutations in IFNγ-pathway genes of moderate or severe variant effect predictor (VEP) impact scoring. Signatures of CD8 T cell and monocytes were retrieved and combined to create a CD8-monocyte signature, from the R package consensusTME(Jiménez-Sánchez et al., 2019), which has curated cell-type signatures for each tumour type. The function geneSetEnrichment was used to perform single-sample gene set enrichment analysis (ssGSEA) for each tumour type using the custom CD8-monocyte signature. For gene signature analysis of publicly available Visium CytAssist spatial transcriptomic datasets, Loupe browser files for human colon adenocarcinoma (FFPE) and human lung squamous cell carcinoma (FFPE) were downloaded from 10X Genomics and analysed using Loupe Browser 7.0.1. Gene sets were retrieved from reference publications and indicated in Supplementary Dataset S1, and visualized as log normalized average expression of all features in the gene set.

### Statistics

Unless otherwise noted, all data involving *in vivo* experiments are pooled from ≥ 2 separate experiments. Statistical analyses were performed using GraphPad Prism software. Error bars represent standard error of mean (SEM) calculated using Prism. Statistical tests used include non-parametric Mann-Whitney *t* tests for comparisons between two groups, and two-way ANOVA using Šídák’s test for multiple comparisons between multiple two or more groups of data.

### Data availability

The mouse scRNAseq data generated in this study have been deposited in the GEO database under accession code GSE260972. Datasets retrieved from 10X Genomics are licensed under the Creative Commons Attribution license. All data are available from the authors upon reasonable requests, as are unique reagents used in this Article.

## Results

### Loss of IFNγ signalling does not result in decreased patient survival for multiple cancer types

While several reports of IFNγ pathway somatic mutations were found in post-ICB treatment patients, it is often unclear whether those mutations can be present before ICB, and if they affect overall survival of cancer patients. We investigated the prevalence of those mutations before ICB by collating data from The Cancer Genome Atlas (TCGA) for pre-treatment tumours which harboured mutations in the IFNγ receptor subunits (*IFNGR1*/*2*) or downstream signalling molecules (*JAK1/2*, *STAT1*). The prevalence of mutations was found to be relatively infrequent (i.e. alteration frequency <10%) across most tumour types except endometrial cancer and melanoma (*Fig.1A*), compared to known oncogenic mutations such as *KRAS* or *PIK3CA* (*Fig.S1A,B*). IFNγ pathway mutations were found at similar frequencies compared to antigen presentation mutations such as in *B2M*, *TAP*, or *HLA* molecules (*Fig.S1C*), suggesting that the pressure induced by T cells is not potent enough in most tumours to select for these types of mutations. More importantly, presence of IFNγ-pathway mutations did not result in significantly higher overall mortality in cancer types with the highest mutational frequency, and even correlated with improved survival in endometrial cancer (*Fig.1B-E*). This, combined with previous preclinical data which showed enhancement of checkpoint blockade responses in pathway mutants, suggests that tumours with mutations in the IFNγ pathway are as efficiently controlled by the immune system as tumours that are sensitive to IFNγ. This led us to investigate the immunological changes in the tumour microenvironment which may be promoting immunity towards these types of tumours.

**Figure 1.**
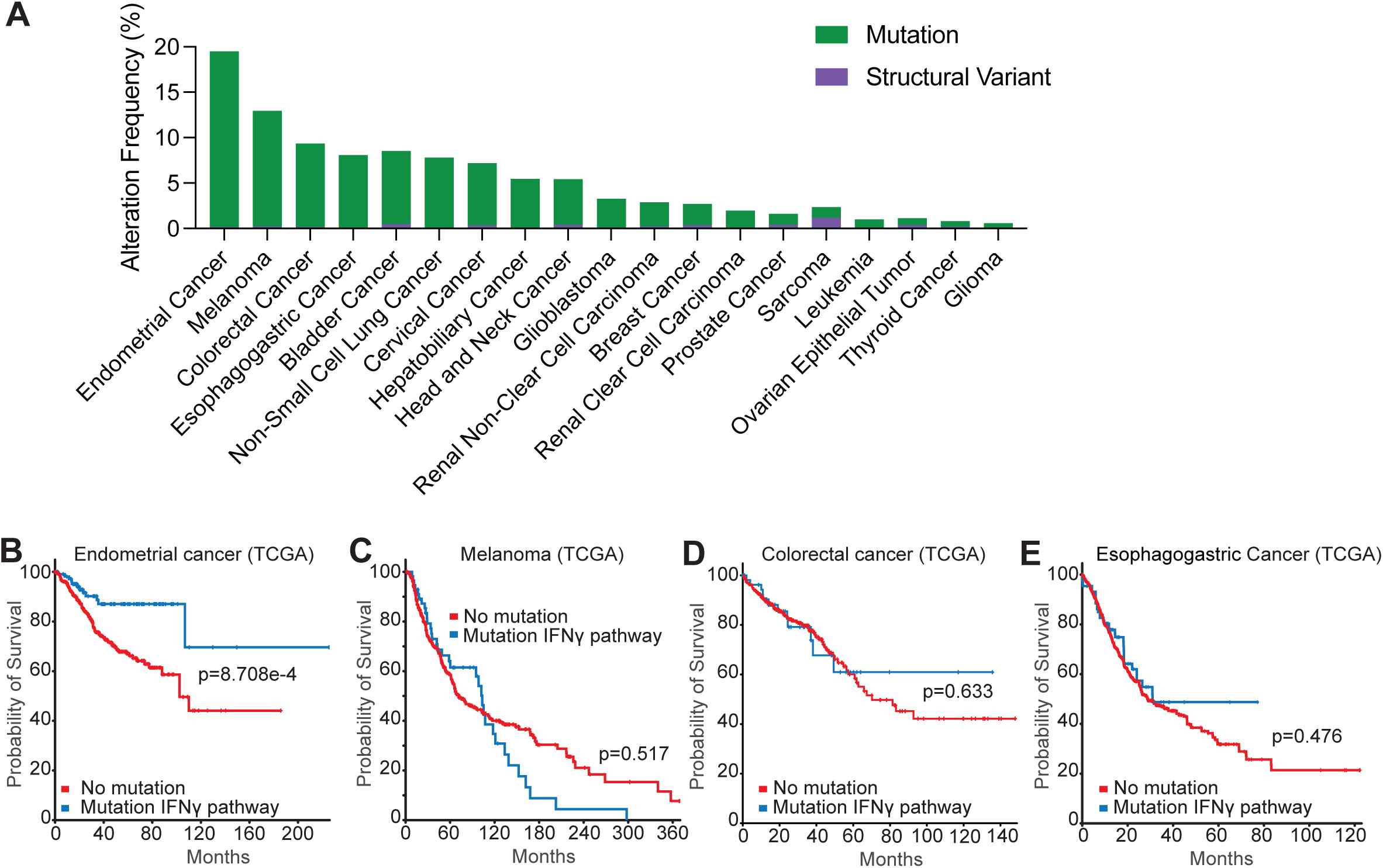
Presence of mutations in IFNγ signalling pathway genes does not preclude decrease in overall survival in clinical data. (**A**) Frequency of alterations in IFNGR1, IFNGR2, JAK1, JAK2, or STAT1 (IFNγ pathway) across cancers in The Cancer Genome Atlas (TCGA), where cases in green represent gene mutations and purple are structural variants of the genes. (**B-E**) Comparison of survival curves of endometrial (**B**, n=462 without and n=111 with mutation in the IFNγ pathway), melanoma (**C**, n=366 without and n=57 with mutation in the IFNγ pathway), colorectal (**D**, n=473 without and n=49 with mutation in the IFNγ pathway), and esophagogastric (**E**, n=1034 without and n=103 with mutation in the IFNγ pathway) human cancers between unaltered cases and cases with mutations in IFNγ pathway genes. *p*-values represent log-rank testing.

### Control of B16F10 IFNγRKO tumours is CD8^+^ T cell-dependent

We established an IFNγ-insensitive model of B16F10 melanoma (*Fig.2A,B*) through CRISPR-Cas9 knockout of IFNγR1 (IFNγRKO). As expected, deletion of IFNγR1 results in lack of MHC class I and II upregulation *in vivo* (*Fig.2C,D)*, and following IFNγ stimulation *in vitro* (*Fig.S2A*). This cell line also expresses ovalbumin (OVA) (B16-OVA) to track antigen/tumour-specific T cells. Given that not all individual cancer cells in human tumours are unresponsive to IFNγ, we used a B16-OVA WT and IFNγRKO admix model whereby tumour cells are tagged in ZsGreen or mCherry, or vice versa, and mixed in equal proportions prior to engraftment (*Fig.2E*). As a control, we admixed ZsGreen and mCherry WT cells (WT:WT) or ZsGreen and mCherry IFNγRKO (KO:KO) cancer cells. No differences in tumour volumes between control WT:WT, KO:KO or admixed WT:KO tumours were observed *in vivo* (*Fig.2F*). However, a phenotype emerged whereby IFNγRKO cells outgrew WT cells in a time-dependent matter (*Fig.2G and S2B*), which we hereafter refer to as selection. This is consistent with human data suggesting that mutations in the IFNγ pathway do not confer any advantage in tumour growth as a whole or survival, but these mutations can rise and take over other clones. To ascertain whether this selection was dependent on IFNγ-signalling by B16 tumour cells, we created B16-OVA cell line expressing a IFNγR1 mutated on Y445A, which abolishes the STAT1 binding site(Greenlund et al., 1995). Similar to complete IFNγR deletion, Y445A mutation resulted in selection, with mutated cells taking over their WT counterparts (*Fig.S2C*). Furthermore, the selection was lost when admixed tumours were engrafted in IFNγKO mice (*Fig.2H*), confirming that selection was induced by IFNγR deletion rather than off-target effects. NK cells have been implicated in the control of tumours with low MHC-I expression(Ljunggren and Kärre, 1990). To test whether NK cells were responsible for controlling IFNγRKO tumours, we treated mice with NK1.1 antibodies to deplete NK cells prior to engraftment of WT or IFNγRKO B16-OVA tumours. Consistent with an early role of NK cells in controlling tumour growth(Stabile et al., 2017), NK depletion led to increased tumour growth and decreased survival (*Fig.S2D,E*). However, IFNγRKO tumour volumes were surprisingly equivalent to that of WT tumours when NK cells were depleted, suggesting that NK cells did not provide enhanced control of tumours with low MHC-I and that other immune cells might be involved in controlling IFNγ-insensitive tumours. B16-OVA IFNγRKO tumours are predicted to be less sensitive to CD8^+^ T cells because of their low expression of MHC-I(Seliger et al., 2001; Yu et al., 2018). To test this, we engrafted either WT, IFNγRKO or admixed B16-OVA in CD8ɑKO mice, which are devoid of CD8^+^ T cells. Selection of IFNγRKO tumour cells was not observed in CD8ɑKO mice; instead, in approximately half the mice, the phenotype was reversed (*Fig.2I*). CD8^+^ T cells are most likely required for selection of IFNγRKO over WT tumours because they preferential target WT tumour cells due to their higher expression of MHC-I(Williams et al., 2020). Unexpectedly, we found tumour growth to be significantly higher in all three tumour types (*Fig.2J*), demonstrating that CD8^+^ T cells are still crucial for controlling IFNγRKO tumours. Consistent with a specific involvement of CD8^+^ T cells in controlling the growth of IFNγRKO tumours, quantification of lymphocyte tumour infiltration by flow cytometry revealed no overt differences except for an increase in OVA-specific CD8^+^ T cells in IFNγRKO and admix tumours (*Fig.2K*), as assessed by OVA-tetramer staining. Thus, despite low MHC-I expression and insensitivity to IFNγ, the control of growth and selection of IFNγRKO tumours remained CD8^+^ T cell- and IFNγ-dependent.

**Figure 2.**
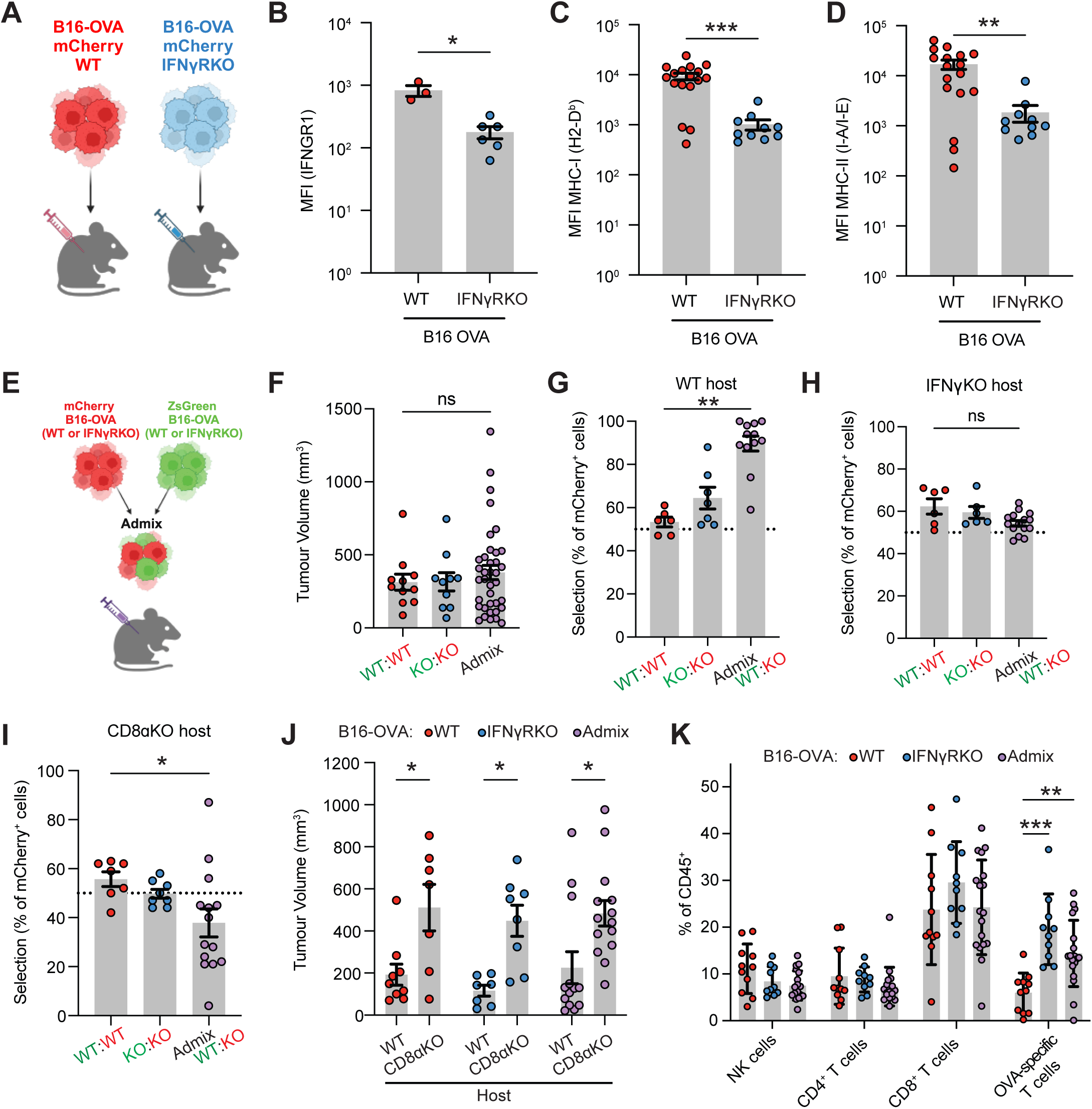
B16F10 melanoma tumours with CRISPR-Cas9 knockout of IFNγR1 are efficiently controlled by the endogenous anti-tumour response. (**A-D**) B16-OVA WT or IFNγRKO cells were engrafted subcutaneously into the flanks of C57Bl/6 WT mice and tumours were harvested after 11-16 days. (**A**) Diagram of the experimental set-up. (**B-D**) Surface expression of IFNγR1 (**B**), MHC-I H2-D^b^ (**C**), and MHC-II I-A/I-E (**D**) expression on mCherry^+^ CD45^-^ cells were analysed by flow cytometry. (**E-H**) WT and IFNγRKO tumours expressing mCherry-OVA or ZsGreen-OVA transgenes were admixed 1:1 prior to engraftment in WT (**F-G**) or IFNγKO (**H**) mice. (**E**) Experimental design. (**F**) Tumour volumes of WT, IFNγRKO, or admixed tumours taken at endpoint on days 12-14 post-engraftment from three independent experiments for WT:WT and KO:KO, and six independent experiments for admixed tumours. (**G-H**) Tumours were harvested and analysed by flow cytometry. Outgrowth of KO tumour cells relative to WT, expressed as percent selection of mCherry^+^ cells in control WT:WT or KO:KO, or WT:KO tumours at day 14-17 in WT (**G**) and IFNγ KO (**H**) mice. Cells were gated on live CD45^-^ mCherry^+^. (**I-J**) WT and IFNγRKO tumours expressing mCherry-OVA or ZsGreen-OVA transgenes were admixed 1:1 prior to engraftment in WT or CD8ɑKO mice. (**I**) Tumours were harvested and analysed by flow cytometry. Graph shows percent selection of mCherry^+^ cells in CD8ɑKO mice. (**J**) Tumour volumes of admixed tumours from WT or CD8ɑKO mice was measured on day 12/13 post-engraftment. (**K**) WT and IFNγRKO tumours expressing mCherry-OVA or ZsGreen-OVA transgenes were admixed 1:1 prior to engraftment in WT mice and harvested at days 12-14 post-engraftment. Infiltration of lymphocyte populations as a percent of total CD45^+^ cells from admixed tumours in WT mice was analysed by flow cytometry. Data are pooled from two or more independent experiments unless otherwise indicated. All data show mean ± SEM with *p*-values by non-parametric Mann-Whitney *t* tests for comparisons between two groups, Kruskal-Wallis tests between three groups with multiple comparisons correction using Dunn’s method, and two-way ANOVA using Šídák’s test for multiple comparisons between multiple two or more groups of data. *p*≤0.05; \*\**p*≤0.01; \*\*\**p*≤0.001.

### scRNAseq identifies significant changes in cytokine signalling and myeloid infiltration of IFNγRKO tumours

Increase in tumour-specific CD8^+^ T cells suggests that inhibition of IFNγ signalling in tumours might alter the cytokine environment, inducing global changes in signalling pathways of other immune subsets which we explored using genomic methods. We employed single cell RNA sequencing (scRNAseq) of the CD45^+^ compartment isolated from pooled tumour samples to better understand the immunological changes which enable effective control of IFNγ-insensitive tumours. Clusters were generated using unsupervised hierarchical clustering and annotated using canonical gene expression patterns (*Fig.3A, S3A*). We identified six myeloid populations. Of those, we found four dendritic cell (DC) subsets: cDC1 (*Xcr1*), cDC2 (*Cd209a*, *Clec10a*), mReg DCs (*Cd200*, *Ccr7*), and plasmacytoid DCs (*Siglech*, *Ccr9*). Other myeloid clusters comprised monocytes (*Ly6c2*, *Ifitm6*, *Vcan*) and macrophages (*C1qa*, *Spp1*). Six lymphoid populations were present in this dataset, which were comprised of NK cells (*Ncr1, Klrb1c*), CD8^+^ T cells (*Cd3, Cd8a, Cd8b1*), regulatory T cells (*Cd4, Foxp3*), cycling T cells (*Mki67, Cd8a*), stem-like T cells (*Cd8a, Tcf7*) and B cells (*Cd19*) (*Fig.3A, S3A*). In comparing the relative frequency of CD45^+^ populations, lymphoid populations were modestly variable between WT and IFNγRKO tumours, whereas the macrophage cluster was expanded in WT compared to IFNγRKO tumours (*Fig.3B*). Gene set enrichment analysis (GSEA) of the entire scRNAseq dataset unexpectedly revealed IFNγ-signalling and related pathways such as antigen presentation as top hits from both hallmark and Reactome databases of immune cells from IFNγRKO tumours (*Fig.3C,D*), suggesting that a highly inflammatory environment emerges following IFNγR deletion in tumour cells. Using ELISA and Legendplex, we confirmed that IFNγRKO tumours contained significantly higher levels of IFNγ and IL-6, whilst other cytokines such as IFNɑ, TNFɑ, M-CSF, IL-4, and IL-10 were similar in concentration (*Fig. 3E-H, S3B-D*), showing that the increase in inflammatory cytokine milieu detected by sequencing in IFNγRKO tumours was the result of an increase in specific inflammatory cytokines. We further employed CellChat as a tool for dissecting soluble signals present in the tumours, as multiple pathways present from GSEA were indicative of intercellular signalling as primary differences between the two tumour types. We found CD8^+^ T cells from IFNγRKO tumours to be main signal receivers from monocytes, whereas mono-macs/macrophages from WT tumours were primary signal senders in WT tumours (*Fig.3I, S3E*). Furthermore, quantifying the differential strength of interactions within soluble signalling pathways showed stronger overall interactions between macrophages and other immune subsets such as Tregs and CD8^+^ T cells in WT tumours compared to IFNγRKO (*Fig.3J*).

**Figure 3.**
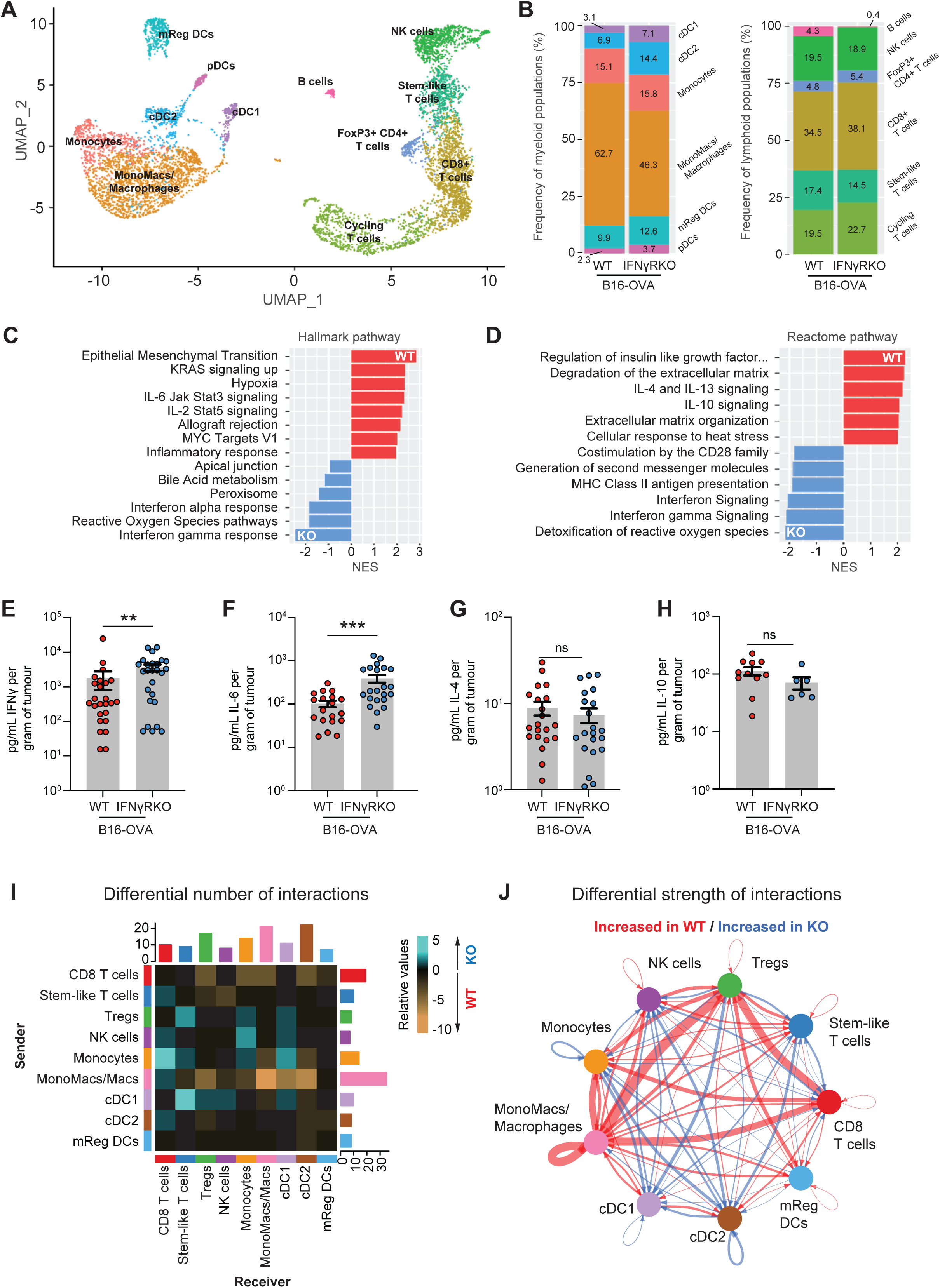
Single-cell RNAseq analysis of CD45^+^ tumour-infiltrating cells reveals the presence of enhanced inflammatory milieu in IFNγRKO tumours. (**A-B**) UMAP projection of CD45^+^ cells and relative abundance of distinct immune populations in WT and IFNγRKO tumours. Clusters show a combined 7014 cells, with 3004 cells from WT tumours, and 4010 cells from IFNγRKO tumours. (**C-D**) Gene set enrichment analysis using hallmark (**C**) or Reactome (**D**) databases for identification of enriched signalling pathways, expressed as normalized enrichment scores (NES). (**E-H**) Intra-tumoural concentrations of IFNγ (**E**), IL-6 (**F**), IL-4 (**G**), IL-10 (**H**) measured by ELISA or LegendPlex using supernatants of *ex vivo* WT (n≤20) and IFNγRKO (n≤20) dissociated tumours, normalized to tumour weight. Data are pooled from four or more independent experiments. Data show mean ± SEM with *p*-values by non-parametric Mann-Whitney *t* tests, \*\**p*≤0.01; \*\*\**p*≤0.001. (**I-J**) CellChat ligand-receptor inference analysis was performed on sc-RNAseq data from (**A**). (**I**) Heatmap of the differential number of interactions between sender (y-axis) and receiver (x-axis) populations. Bar plots on each axis represents the sum of all interactions in absolute values for each sender or receiver cell type. (**J**) Circle plot visualizing strength of signalling interactions between immune populations from WT and IFNγRKO tumours. Vertices represent independent populations, and arrows indicate direction of signals sent, where broader lines represent increased communication probability of signalling interactions.

Overall, our data indicate that IFNγR deletion in tumour cells triggers a remodelling of the immune response and mediators centred around monocytes/macrophages. This led to our hypothesis that tumour-infiltrating monocytes and tumour-associated macrophages may be key in modulating lymphocyte function in each tumour microenvironment.

### Inflammatory myeloid subsets are enhanced by the tumour microenvironment of IFNγ-insensitive tumours

Following the observation that differences in cytokine signalling and CD45 subsets were likely to lie within the myeloid population, we subset monocytes and macrophages and identified six clusters with unique gene signatures *(Fig.4A and S4A)*. In line with recent scRNAseq studies describing murine tumour-infiltrating myeloid cells populations(Zhang et al., 2020; Zilionis et al., 2019),(Mulder et al., 2021), we identified three monocytic populations, namely non-classical monocytes (Cluster 6; *Nr4a1, Ifitm6*), tumour-infiltrating monocytes (Cluster 2; *Ly6c2*, *Vcan*), and transitionary mono-mac (Cluster 3; *Adgre1*, *Folr2*). We also described four tumour-associated macrophage (TAM) populations, namely IFN-stimulated TAMs (Cluster 1; *Nos2, Sod2*), regulatory TAMs (Cluster 0; *Arg1*, *Spp1*), angiogenic TAMs (Cluster 5; *Vegfa*), and complement TAMs (Cluster 4; *C1qa*) (*Fig.4B*). IFNγRKO tumours were dominated by non-classical and tumour-infiltrating monocyte clusters, and IFN-stimulated TAMs, whereas regulatory TAMs were unique to WT tumours (*Fig.4C*). Trajectory analysis supported the branching of pre-macrophage (cluster 3) into more differentiated macrophage phenotypes, i.e. regulatory TAMs (cluster 0), IFN-stimulated TAMs (cluster 1), and angiogenic TAMs (cluster 5) (*Fig.4D*). This suggests that the macrophages present in our model might partially be derived from monocytes. Those monocyte-derived macrophages are known to accumulate over time during tumour progression, and have been suggested to shape immune responses(Casanova-Acebes et al., 2021; Franklin et al., 2014). However, the type of macrophages elicited in WT versus IFNγRKO tumours differs. Indeed, module scoring of angiogenic and regulatory TAM signatures, taken from recent literature(Ma et al., 2022), confirmed the overall increased presence of these subsets in WT tumours compared to IFNγRKO (*Fig.4E*, *Table 1*). Angiogenic and regulatory TAMs are known to be pro-tumourigenic(Ma et al., 2022; Mantovani et al., 2017), and we hypothesise that their absence in IFNγRKO tumours might contribute to tumour control.

**Figure 4.**
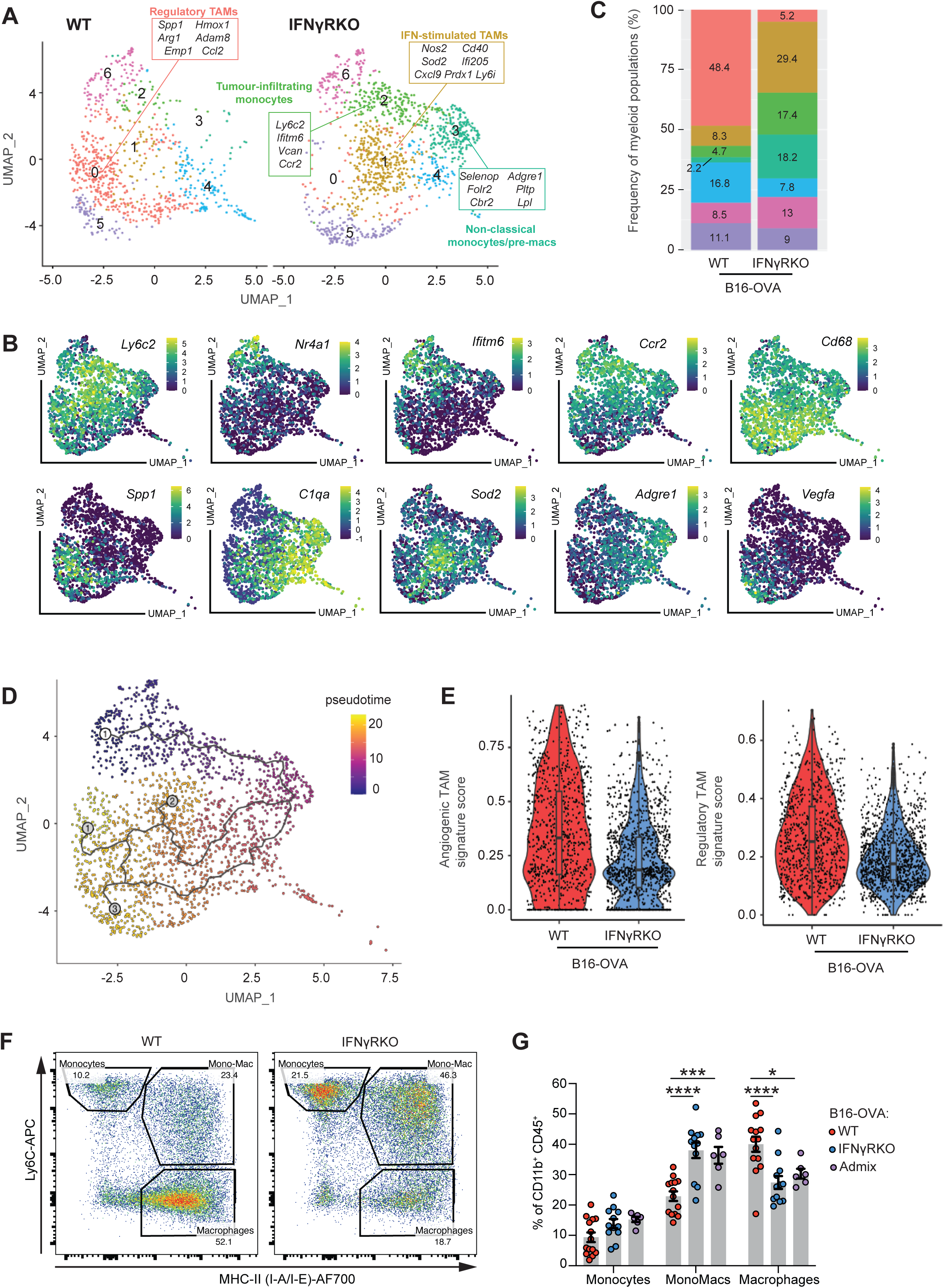
Less differentiated monocyte-macrophage subsets with pro-inflammatory signatures are prominent in IFNγRKO tumours. (**A-E**) Myeloid cells from the scRNA-seq data from Fig.3A were subset and re-clustered. (**A**) UMAP of WT and IFNγRKO monocyte/macrophage subclusters with distinct myeloid subtypes highlighted by representative gene signatures. (**B**) Feature plots showing relative gene expression of key monocyte/macrophage genes. (**C**) Relative frequencies of each subcluster expressed as stacked bar plots. (**D**) Trajectory analysis overlaid on the UMAP projection of monocyte/macrophage cell clusters, coloured by pseudotime. (**E**) Violin plots comparing module scoring of angiogenic and regulatory TAMs gene signatures of macrophage subclusters of WT and IFNγRKO tumour samples. (**F-G**) WT, IFNγRKO or admix tumours were engrafted in WT mice and analysed by flow cytometry after 14 days. (**F**) Representative flow plots of tumour-infiltrating CD45^+^ CD11b^+^ cells from WT and IFNγRKO tumours, gated by Ly6C and MHC-II expression for delineation of monocyte, mono-mac and macrophage populations. Plots are representative of three or more independent experiments. (**G**) Relative frequencies of each gated population from WT, IFNγRKO or admixed tumours. Data are pooled from three independent experiments. Data show mean ± SEM with *p*-values by two-way ANOVA using Šídák’s test for multiple comparisons, *p*≤0.05; \*\**p*≤0.01; \*\*\**p*≤0.001, \*\*\*\**p*≤0.0001.

**Table 1.**
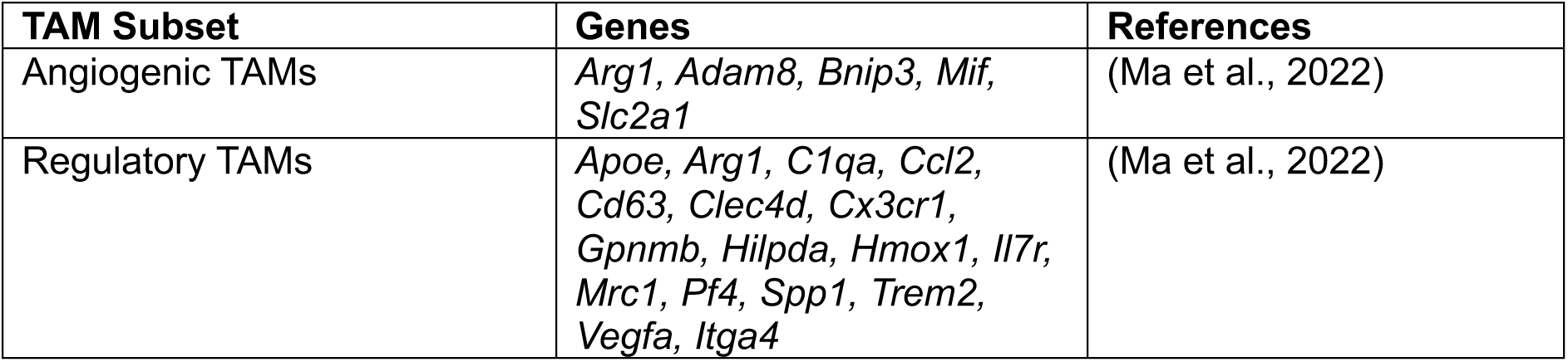
Gene signatures for module and enrichment scoring.

We then sought to confirm our scRNAseq findings using flow cytometry and classified monocytes, mono-macs, and macrophages according to Ly6C and MHC class II expression, with gated populations corresponding well to co-expression of macrophage markers such as F4/80, CD204, CD206 and TREM2 (*Fig.4F, S4B,C*). Using this gating strategy to delineate monocytes, mono-macs and macrophages, IFNγRKO and admixed tumours retained a significant proportion of myeloid cells which were monocytic in origin compared to WT (*Fig.4G*). We then used spectral flow cytometry for deeper phenotyping of the myeloid landscape (*Table 2*). Unsupervised clustering of spectral flow cytometry data of the CD11b^+^ CD45^+^ population revealed an increase in clusters 1 and 3 in IFNγRKO tumours, which represent a population of monocytes characterised by Ly6C^hi^ CD86^+^ or CD62L^+^ (*Fig.S4D-F*). In addition, cluster 7 was increased in WT tumours and corresponded to F4/80^+^ and CD68^+^ population, indicative of a more differentiated macrophage phenotype.

**Table 2.**
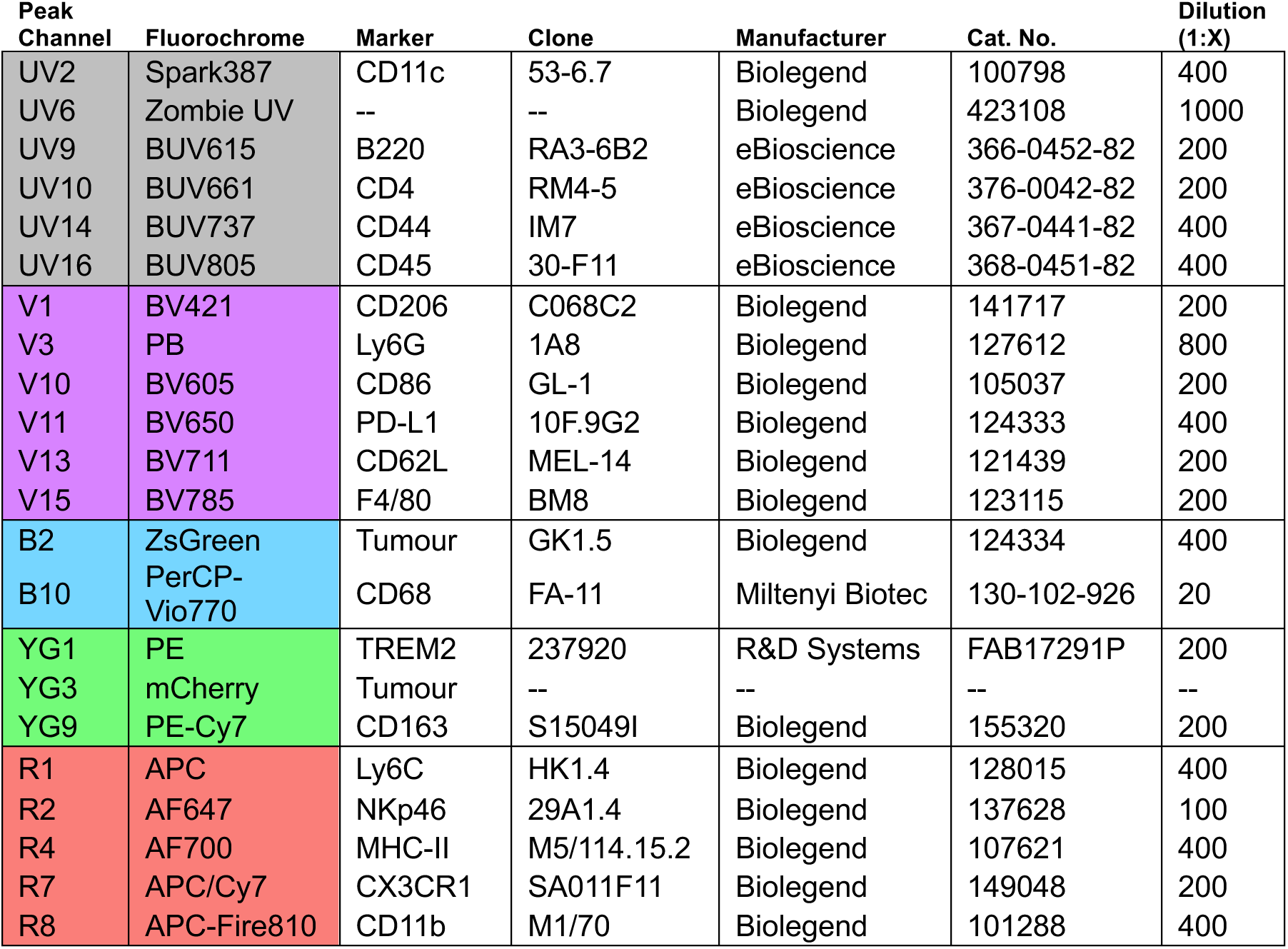
Spectral Flow Cytometry Markers.

We concluded that IFNγR deletion in tumour cells induces a remodelling of the myeloid compartments, with an increase in monocytes and inflammatory macrophages and a concomitant diminution of regulatory TAM.

### Monocyte recruitment is required for controlling IFNγ-insensitive tumours

The CCL2-CCR2 signalling axis plays an indispensable role in recruitment and trafficking of myeloid populations during infection and inflammation(Kurihara et al., 1997; Si et al., 2010). Given that Ly6C^hi^ inflammatory monocytes primarily depend on CCR2 for tumour infiltration(Qian et al., 2011) and TAMs originate from recruited monocytes as well as tissue-resident macrophages(Franklin et al., 2014), we engrafted B16-OVA WT or IFNγRKO tumours into CCR2KO mice to determine whether impeding CCR2-dependent recruitment would impact tumour growth. Monocyte recruitment was almost entirely impeded in CCR2KO tumours, which also significantly lacked mono-mac and macrophage populations, indicating that monocytes in this model were indispensable for macrophage recruitment and/or differentiation (*Fig.5A*). As observed in previous tumour studies using CCR2KO mice(Sawanobori et al., 2008), monocyte and macrophage populations are replaced by granulocytes, which we observed as a neutrophilic influx especially in IFNγRKO tumours. Importantly, IFNγRKO tumours grew significantly faster than WT when engrafted into CCR2KO mice (*Fig.5B*), suggesting that monocyte recruitment is required for the control of IFNγRKO tumours. Deletion of CCR2 resulted in CD8^+^ T cell and NK cell expansion in WT, but not IFNγRKO, tumours (*Fig.5C*), suggesting that loss of phenotypically immunosuppressive macrophages in WT tumours (*Fig.4A-B*) aids in unleashing lymphocyte activity, which in turn improves WT tumour control(Kersten et al., 2022; Park et al., 2023). Loss of CCR2 also decreased the frequency of OVA-specific T cells in IFNγRKO-tumour bearing mice compared with WT mice (*Fig.5D*), which indicates that monocyte recruitment is necessary for the recruitment and/or retention of antigen/tumour-specific CD8^+^ T cells in IFNγ-insensitive tumours.

**Figure 5.**
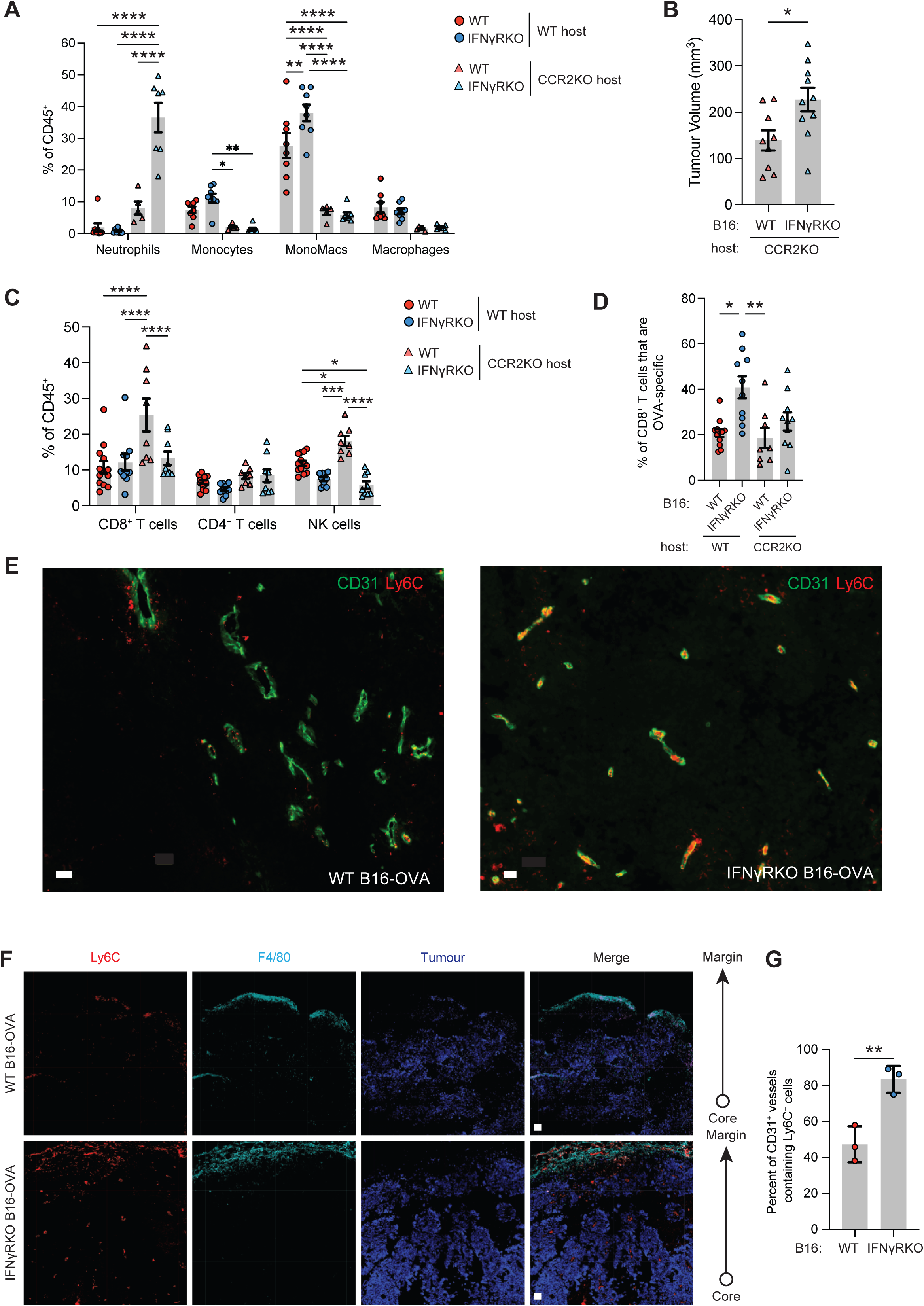
Control of IFNγRKO tumours is diminished in CCR2KO mice following lack of monocyte recruitment. (**A-D**) WT, IFNγRKO or admix tumours were engrafted in WT or CCR2KO mice. (**A**) Infiltration of myeloid populations relative to total CD45^+^ cells in WT or IFNγRKO tumours engrafted into WT or CCR2KO mice. (**B**) Tumour volumes of WT or IFNγRKO tumours measured at endpoint on day 13 post-engraftment. (**C**) Infiltration of T cells and NK cells relative to total CD45^+^ cells. (**D**) Frequency of OVA-specific T cells as a proportion of CD8^+^ T cells. Data are from two independent experiments. (**E-G**) Frozen sections from WT or IFNγRKO tumours engrafted in WT mice were stained with the indicated markers and imaged. (**E-F**) Representative images indicating location of Ly6C (red) and F4/80 (Cyan) expressing cells relative to CD31^+^ (green) blood vessels. Scale bar = 40um (**E**), and 100um (**F**). (**G**) Graph shows the percentage of blood vessels that are in contact with Ly6C^+^ cells. Each dot is a tumour (n=3). All data show mean ± SEM with *p*-values by non-parametric Mann-Whitney *t* tests for comparisons between two groups, and two-way ANOVA using Šídák’s test for multiple comparisons between multiple two or more groups of data. *p*≤0.05; \*\**p*≤0.01; \*\*\**p*≤0.001, \*\*\*\**p*≤0.0001.

Consistent with the recruitment of monocytes from the periphery, imaging of WT and IFNγRKO tumours revealed that Ly6C^+^ cells resided in or near CD31^+^ blood vessels, which were found either within the core of the tumour, or at the margin, surrounding the tumour (*Fig.5E, S5A,B*). Interestingly, Ly6C^+^ cells located at the margin also expressed F4/80, suggesting that myeloid cells become excluded from the tumour core as they differentiate (*Fig.5F, S5B*). WT and IFNγRKO tumours display gross similar location of the myeloid subsets, showing that the spatial distribution of myeloid cells was not impacted by IFNγR deletion in tumour cells. Focusing on the core of the tumour to specifically investigate the relationship between monocytes and blood vessels, we observed an increased presence of monocytes in contact with blood vessels in IFNγRKO compared to WT tumours, in agreement with enhanced recruitment of monocytes in IFNγRKO tumours (*Fig.5G*).

Overall, our results indicate that monocyte infiltration can promote myeloid-CD8^+^ T cell crosstalk which is important for controlling IFNγ-insensitive tumours. As IFNγRKO tumour control is lost in CCR2KO, modulation of these populations rather than complete inhibition of recruitment would be a more compelling strategy for altering CD8^+^ T cell function.

### IFNγ-insensitive tumours support a pro-inflammatory microenvironment driven by CD8^+^ T cells which promotes monocyte infiltration and mono-mac differentiation

Our scRNAseq data analysis indicated an increase in the inflammatory milieu following IFNγR deletion. Because the tumours themselves were no longer sensitive to IFNγ, we reasoned that IFNγ was acting on other cells. To explore this, we scored IFNγ signalling in our transcriptomics dataset using a well-established IFNγ gene signature(Ayers et al., 2017). IFNγR deletion in tumours led to an increase in IFNγ signalling primarily in the myeloid compartment (*Fig.6A*), suggesting that IFNγ was important for the remodelling of the myeloid compartment in IFNγRKO tumours. Indeed, the increase in monocyte and mono-macs observed in IFNγRKO tumours (*Fig.4G*) was lost if those tumours were engrafted in IFNγKO mice (*Fig.6B*). In addition, both WT and IFNγRKO tumours were no longer controlled when engrafted in IFNγKO mice (*Fig.6C*), demonstrating that IFNγ plays an equally important role for both tumour microenvironments. Given the importance of IFNγ for controlling IFNγRKO tumours and remodelling of the myeloid landscape, we sought to identify the cellular source of IFNγ in WT and IFNγRKO tumours. To do so, we used a reporter mouse for IFNγ where EYFP expression is controlled by the IFNγ promoter(Reinhardt et al., 2009). Both CD8^+^ T cells and NK cells were found to be the primary producers of IFNγ in both WT and IFNγRKO tumours during earlier stages of tumour development (i.e. day 7-8 post-engraftment) (*Fig.6D,E*). IFNγ production peaked approximately 10 days post-tumour induction (*Fig.S6A*) and was overtaken by CD8^+^ T cells during latter stages of tumour progression (i.e. days 14-16), regardless of the tumour type. Thus, NK cells lose their capacity to produce IFNγ over time and as such, CD8^+^ T cells remain the main source of IFNγ, even in IFNγRKO tumours.

**Figure 6.**
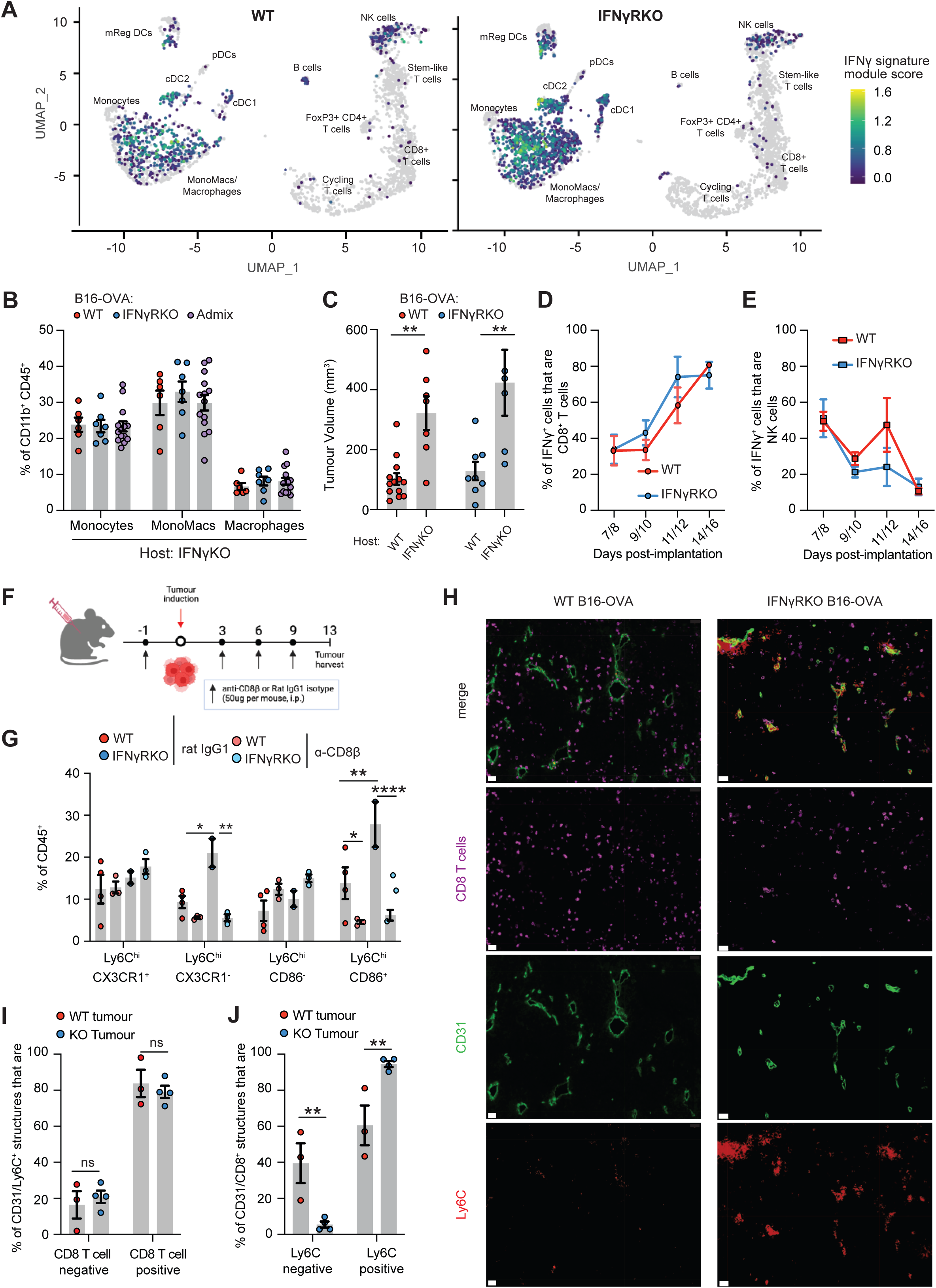
Recruitment of monocytes is driven by CD8^+^ T cell-derived IFNγ. (**A**) Module scoring of an IFNγ gene signature on the single-cell data set from Fig.3A. (**B**) Infiltration of myeloid populations relative to total CD11b^+^CD45^+^ cells in WT, IFNγRKO or admixed tumours engrafted into IFNγKO mice. Data are pooled from two independent experiments. (**C**) Tumour volumes of WT or IFNγRKO tumours measured on day 10 post-engraftment of WT or IFNγKO mice. Data are pooled from two independent experiments. (**D-E**) WT or IFNγRKO tumours were engrafted in GREAT mice and harvested when indicated. Percentage of cells which are IFNγ+, as measured by EYFP expression by tumour-infiltrating CD8^+^ T cells (**D**) and NK cells (**E**). Data are pooled from four independent experiments, with timepoints varying between experiments. (**F-G**) Antibody depletion of CD8^+^ T cells using anti-CD8β prior to and following tumour engraftment. (**F**) Experimental design. (**G**) Frequency of specific Ly6C^hi^ monocyte subsets following CD8^+^ T cell depletion was analysed by flow cytometry 13 days post-engraftment. (**H-J**) Frozen sections from WT or IFNγRKO tumours were stained with the indicated markers and imaged. (**H**) Representative immunofluorescence images taken at the core of WT and IFNγRKO tumours showing the location of Ly6C (red) and CD8 (magenta) cells relative to CD31^+^ blood vessels (green). Scale bar = 30um. (**I**) Bar graph shows the percentage of CD31^+^/Ly6C^+^ structures that contain CD8^+^ T cells. (**J**) Bar graph shows the percentage of CD31^+^/CD8^+^ T cell structures that contain Ly6C cells. Each dot is a tumour (n=3-4). All data show mean ± SEM with Kruskal-Wallis testing between three groups with multiple comparisons correction using Dunn’s method, and two-way ANOVA using Šídák’s test for multiple comparisons between multiple two or more groups of data. * *p*≤0.05; \*\**p*≤0.01; \*\*\*\**p*≤0.0001.

As CD8^+^ T cells were important for limiting IFNγRKO tumour growth (*Fig.2B*), and the production of IFNγ enables the inflammatory myeloid landscape responsible for IFNγRKO tumour control, we hypothesised that CD8^+^ T cells regulate the myeloid landscape in IFNγ-insensitive tumours. As for IFNγKO mice, we did not observe an increase in monocytes and mono-macs in IFNγRKO compared to WT tumours when engrafted in CD8ɑKO mice (*Fig.S6B*), indicating that CD8^+^ T cells were important to recruit monocytes to IFNγRKO tumours. Similar data were obtained when we specifically depleted CD8^+^ T cells by treating mice with a CD8ß depleting antibody before implanting WT or IFNγRKO tumours (*Fig.6F*). Deeper phenotyping revealed that loss of CD8^+^ T cells resulted in depletion of classical monocyte Ly6C^hi^ CX3CR1^-^ or CD86^+^ subsets in both in WT and IFNγRKO tumours (*Fig.6G*), but did not affect F4/80^+^ macrophage populations (*Fig.S6C-E*). Ly6C^hi^ CX3CR1^-^ monocytes have previously been described for their role in renewing intra-tumoural TAM populations(Movahedi et al., 2010). This overall indicates that CD8^+^ T cells are required for the recruitment of classical monocytes with inflammatory and costimulatory properties.

In agreement with the role of CD8^+^ T cells in recruiting monocytes, we observed that most blood vessels, labelled with CD31, were surrounded by CD8^+^ T cells (*Fig.6H, S6F,G*). At the margin, Ly6C^+^ monocytes were associated with those structures in both WT and IFNγRKO tumours (*Fig.S6F,G*), but due to the large presence of differentiated, F4/80^+^, myeloid cells at the margin, it was unclear whether this could be associated with active recruitment. We therefore focused on blood vessels present in the core of the tumour for quantification. To infer whether the presence of monocytes in or close to vessels correlated with the presence of CD8^+^ T cells, we focused on vessels that contained Ly6C^+^ cells and quantified the proportion of those structures that included CD8^+^ T cells. More than 80% of vessels that contained Ly6C^+^ cells also contained CD8^+^ T cells, regardless of the tumour type, confirming that CD8^+^ T cells are required for monocyte recruitment in both WT and IFNγRKO tumours (*Fig.6I*). To understand whether CD8^+^ T cells had the same ability to recruit monocytes in WT and IFNγRKO tumours, we focused on vessels that were in contact with CD8^+^ T cells and quantified the proportion of those structures that contained Ly6C^+^ cells. While almost all CD8/vessel structures contained Ly6C^+^ cells in IFNγRKO tumours, only 60% did so in WT tumours (*Fig.6H,J*), consistent with the CD8^+^ T-cell-dependent increase in monocyte recruitment observed in IFNγRKO tumours. Thus, while monocytes need CD8^+^ T cells for their recruitment regardless of the tumour type, inhibition of IFNγ sensing in tumours leads to an increase in the ability of CD8^+^ T cells to recruit monocytes. These findings are consistent with studies which describe the role of pro-inflammatory cytokines such as IFNγ in enabling leukocyte adhesion and transendothelial migration through integrin(Carman and Martinelli, 2015; Zhang et al., 2011) or MHC class II upregulation(Masuyama et al., 1986). In IFNγRKO tumours, higher IFNγ levels may increase adhesion of lymphocytes and monocytes to intra-tumoural endothelium, which we observe as a quantifiable increase in these cell-cell interactions.

Overall, our data demonstrate that a CD8/monocyte cross-talk is potentiated in tumours insensitive to IFNγ and underlies their control.

### Elevated CD8-monocyte immune signature scoring across multiple human cancer types

Our data using mouse models points to a remodelling of the immune response driven by inhibition of IFNγ receptor or signalling. To explore whether this is also elicited in human tumours, we elected to investigate whether enrichment scoring using a combined CD8-monocyte signature would be increased in TCGA RNAseq datasets in which the patient tumours harboured mutations in *IFNGR1/2*, *JAK1/2* or *STAT1*. Only datasets in which the calculated variant consequences (i.e. VEP IMPACT score) deemed high or moderate were included for analysis. Using single-sample gene set enrichment analysis, multiple tumour types scored higher for CD8-monocyte enrichment in IFNγ-pathway mutation-containing datasets compared to controls (*Fig.7A*). This is consistent with our data in mice, and suggests that mutations in the IFNγ-pathway can drive an enhanced CD8^+^ T cell/monocyte immune response.

**Figure 7.**
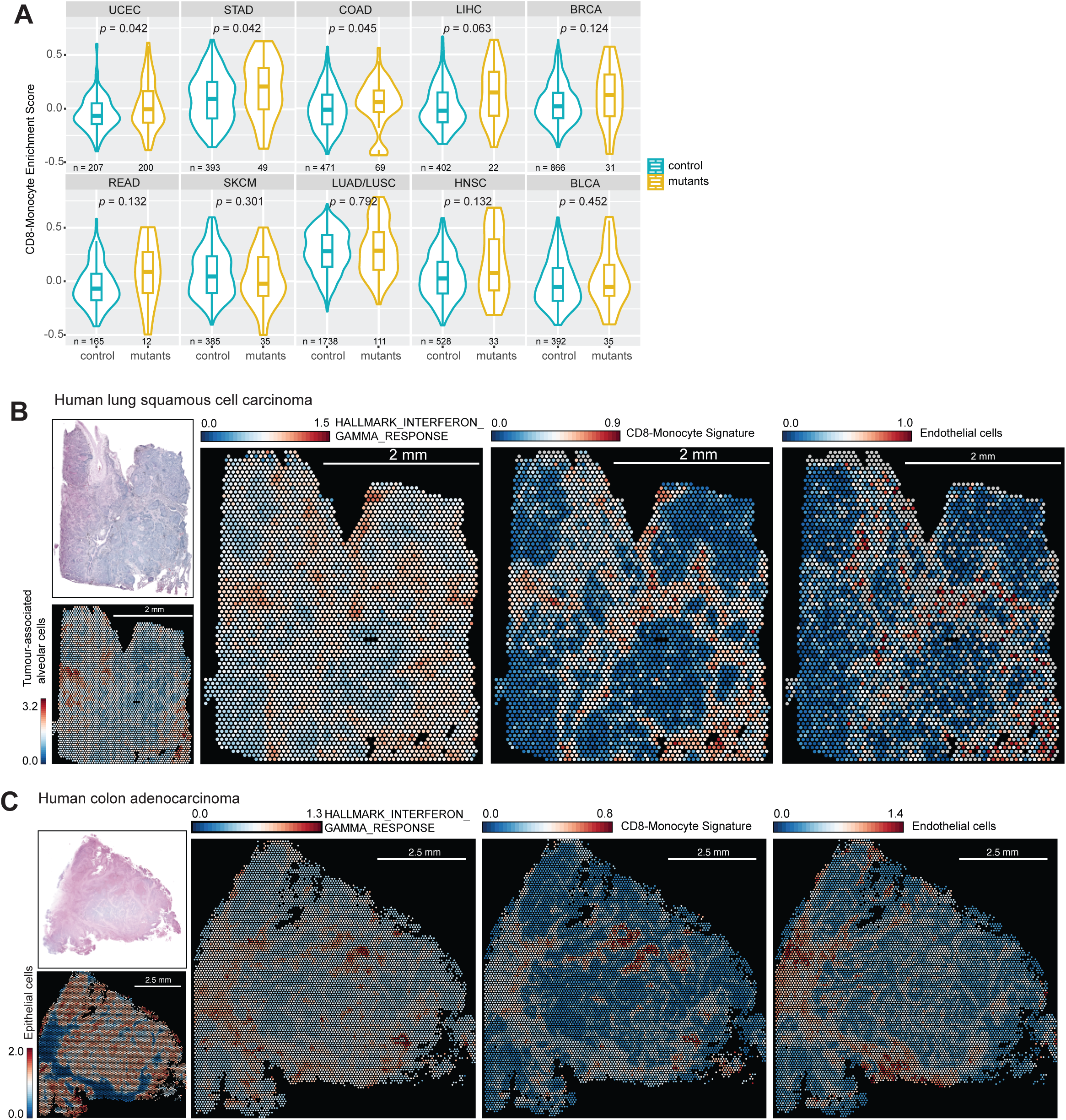
CD8-monocyte signatures are elevated in human tumours with identified IFNγ-pathway mutations, and spatially overlap with IFNγ response signatures. (**A**) Enrichment scoring of CD8-monocyte gene signatures using single gene set enrichment analysis of human TCGA RNAseq datasets with and without IFNγ-pathway mutations. Normalized gene counts from each tumour type were taken from samples which had moderate or high impact *IFNGR1/2*, *JAK1/2*, *STAT1* mutations determined by whole-exome sequencing. Number of samples included for analysis are indicated for each sample set, and statistical testing using Wilcoxon signed-rank test with adjust p-values by false discovery rate testing is shown. (**B**) Analysis of 10X Genomics Visium datasets for hallmark IFNγ response, CD8-monocyte, and endothelial cell gene signatures for human lung squamous cell carcinoma (**B**) or colon adenocarcinoma (**C**) samples. Gene set expression is indicated by heatmap, where colours represent log-normalized average expression.

Our data suggested that the interplay between CD8^+^ T cells and myeloid cells occurred around blood vessels. To explore this in human tumours, we performed analysis of publicly available spatial transcriptomic datasets from the 10X Genomics Visium platform to investigate IFNγ response and CD8-monocyte gene signatures on human lung and colon cancer samples. Across both tumour types, CD8-monocyte gene signature expression coincided spatially with hallmark IFNγ response signatures and endothelial cell markers (*Fig.7B,C, Fig.S7A,B*). These regions were also correlated with M2 macrophage signatures, indicating that IFNγ-response regions are likely associated with active immune infiltration rather than specific immune subsets due to limits in spatial resolution of individual cell types (*Fig.S7C,D*). Other hallmark pathways such as TGFβ-signalling or hypoxia showed incongruent overlap with immune-rich regions, providing evidence that regions of opposing immune function or activity are likely to be compartmentalized.

Overall, analysis of human datasets corresponds to our murine model, and shows that CD8-monocyte signatures can be found in tumours with mutations in the IFNγ-pathway. Human spatial datasets also mirror the association of CD8^+^ T cells and monocytes in our imaging studies, highlighting the importance of their co-operativity in the tumour microenvironment.

## Discussion

In this study, we demonstrate a novel mechanism in which IFNγ-insensitive tumours trigger re-modelling of the tumour microenvironment through accumulation of IFNγ which leads to effective tumour control. We show that IFNγRKO tumours were dominated by inflammatory monocytic and pre-macrophage subsets compared to archetypal tumour-associated macrophages in WT tumours, and this mechanism relied on CCR2-dependent myeloid recruitment. Moreover, we uncovered a monocyte-CD8^+^ T cell reciprocity where depletion of either monocytes or CD8^+^ T cells impaired control of IFNγRKO tumours; loss of monocyte infiltration impeded infiltration of tumour-specific T cells, and CD8^+^ T cell depletion resulted in loss of inflammatory monocyte subsets. The phenomenon of immune sensitization following IFNγ-signalling ablation is not restricted to B16F10 melanoma, as other groups have reported similar findings in mammary, colon, pancreatic and lung tumour models in both Balb/c and C57Bl/6 animals(Dubrot et al., 2022; Lawson et al., 2020; Wang et al., 2021). Incidentally, our results provided a mechanistic basis for reports which showed that IFNγ-pathway mutations undergo positive selection during *in vivo* CRISPR screens(Lawson et al., 2020), and perhaps why mutations in the IFNγ pathway resulted in sensitization towards ICB responses(Dubrot et al., 2022).

We demonstrate that low, baseline levels of MHC class I can be sufficient for eliciting strong CD8^+^ T cell-dependent anti-tumour activity. Consistent with this, meta-analysis studies failed to demonstrate a strong association between stable/progressive disease and loss of HLA(Litchfield et al., 2021). In addition, MHC-I negative tumours only indicate poor survival when PD-L1 is concomitantly expressed, and tumours which are negative for both show no difference in survival^57^. Finally, patient-derived melanoma cell lines carrying JAK1 or JAK2 knockouts retained basal MHC-I expression and the capacity to activate tumour-specific T cells *in vitro*(Torrejon et al., 2020). In this context, our study strongly suggests that IFNγ likely plays a significant role in stabilizing antigen presentation by myeloid subsets, especially during early tumourigenesis, which in turn modulates long-term tumour-specific T cell persistence. We and others have recently highlighted the importance of such myeloid-T cell interactions(Kersten et al., 2022) and other studies have shown that macrophages are capable of cross-presenting tumour antigens to CD8^+^ T cells(Barrio et al., 2012; Sheng et al., 2017).

Although it is often assumed that the most important function of CD8^+^ T cells is direct lysis of the tumour, our data highlights that their role in promoting a cytokine environment permissive for tumour control is equally as important. CD8^+^ T cell-derived IFNγ is known for mobilizing rapid effector functions of innate populations during secondary recall responses during infection, and myeloid cells lacking IFNγR expression fail to control pathogens(Soudja et al., 2014). In ICB-sensitive models such as MC38 murine colon adenocarcinoma, anti-PD-L1 drove a significantly more pro-inflammatory macrophage phenotype compared to untreated tumours, and IFNγR^-/-^ bone marrow transferred into tumour-bearing WT mice were only able to produce M2-like macrophages(Xiong et al., 2019). Accordingly, analysis of patient outcomes following ICB +/-chemotherapy(Carroll et al., 2023; Hammerl et al., 2021; Hwang et al., 2020; Yang et al., 2023) or adoptive cell therapy(Barras et al., 2024) demonstrates that the presence of inflammatory monocytes and M1-like macrophages are substantially better in predicting therapy response than traditional biomarkers such as tumour mutational burden or PD-1 expression(McGrail et al., 2021; Placke et al., 2023; Strickler et al., 2021).

One major question remaining is how these plastic and transitionary myeloid populations shift during disease progression, and whether remodelling of the anti-tumour response also occurs early in human cancers. Sampling of early-stage tumours suggested that tumour CD14^+^ cells were not primarily immunosuppressive against T cell cytokine production or proliferation(Singhal et al., 2019). Our admixed model suggests that immune-mediated clonal selection occurs over a longer period of time, where WT and IFNγRKO tumour cells no longer face equal pressure by CD8^+^ T cells, and loss of the ability to kill MHC-I^low^ cells may be attributed to an increasingly immunosuppressive myeloid compartment. Dissecting out the influence of myeloid cells on CD8^+^ T cell function in a longitudinal manner would assist in answering this fundamental mechanism of escape and selection relative to human clinical scenarios where tumours are heterogeneous in their mutations.

Finally, a monocyte/CD8 T cell signature has been recently described in multiple studies as a predictor of good clinical response following multiple types of immunotherapies(Carroll et al., 2023; Padgett et al., 2023). Our data demonstrate that monocyte/CD8 T cell crosstalk indeed enhances tumour control, in particular for tumours that are less immunogenic due to inhibition of the IFNγ pathway and low MHC expression, and importantly, gives a potential mechanism explaining how this crosstalk potentiates anti-tumour responses, beyond IFNγ-insensitive tumours.

## Acknowledgements

We would like to thank the Wellcome Trust Centre for Human Genetics for the generation of the sequencing data, J. Webber for the assistance with cell sorting, and the dynamic platform and microscopy facility at the Kennedy Institute. This work was supported by Cancer Research UK (CR-UK) (C5255/A18085 through the Cancer Research UK Oxford Centre and 29549 to A.G); the Kennedy Trust for Rheumatology Research (KENN151607 and KENN202112 to A.G), John Fell Funds (0013739 to A.G), and Wellcome Trust studentship and Clarendon Scholarship (V.W.C.L).

## Author contributions

V.W.C.L. and A.G. designed the experiments, analysed the data, and wrote the manuscript. V.W.C.L., G.M., J.M.M. and A.K. performed *in vivo* experiments and processed samples for analysis. V.W.C.L. performed bioinformatic analysis of sequencing and Visium datasets. G.M. prepared imaging samples and performed imaging data collection. G.M. and A.G. performed imaging data analysis. V.W.C.L. and A.G. analysed human data sets. E.W.R. provided reagents and technical advice for experiments. V.W.C.L., U.G., G.P., V.C., and A.G. contributed to the initial conception of the study.

## Supplementary data

**Figure S1.**
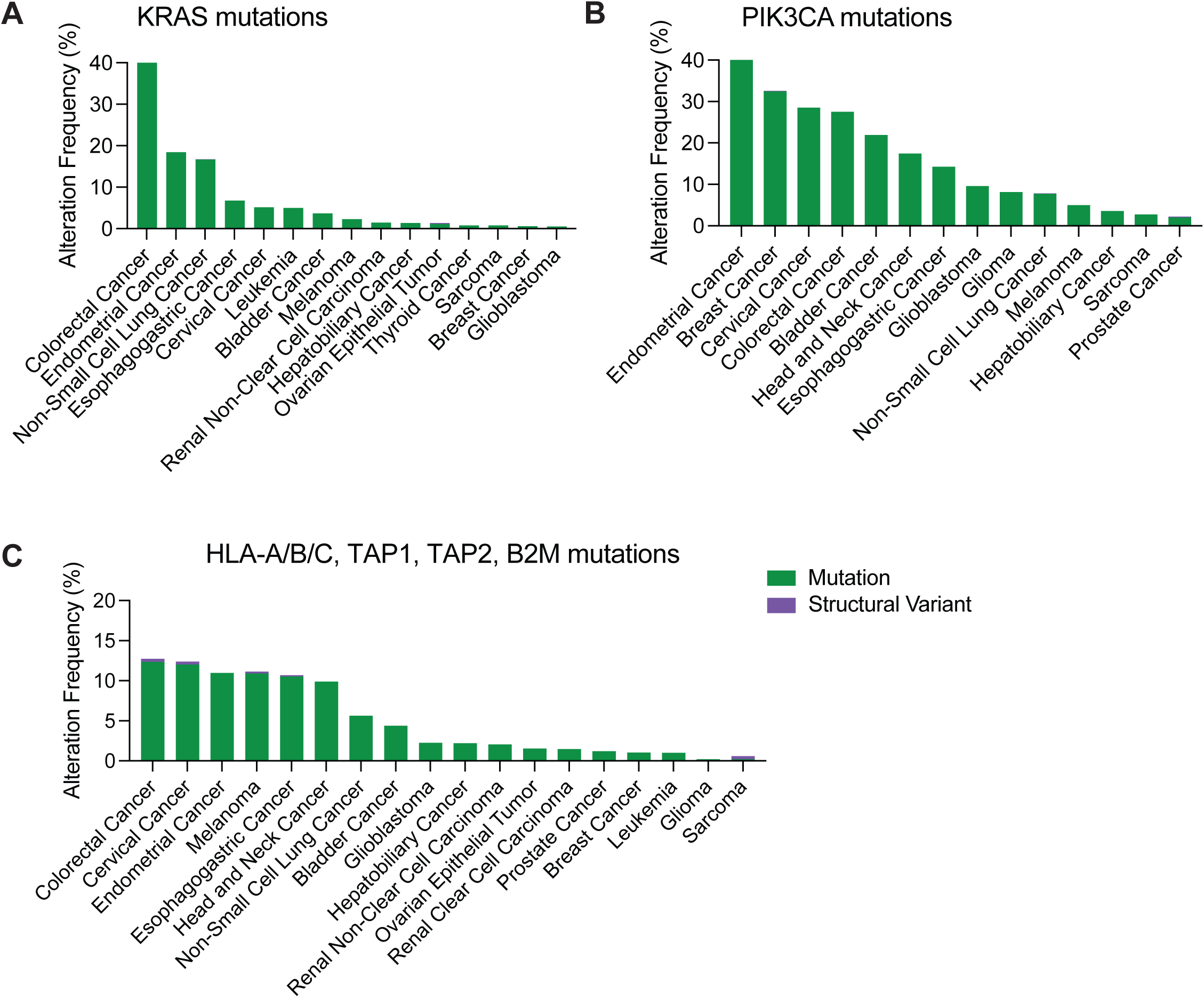
Relative frequency of mutations in different cancer types. Relative frequency of KRAS (**A**), PIK3CA (**B**), and HLA-pathway (**C**) mutations in human cancers from cBioPortal, shown as a percentage of all cases in the database. Mutations in genes are shown in green bars and structural variants in purple.

**Figure S2.**
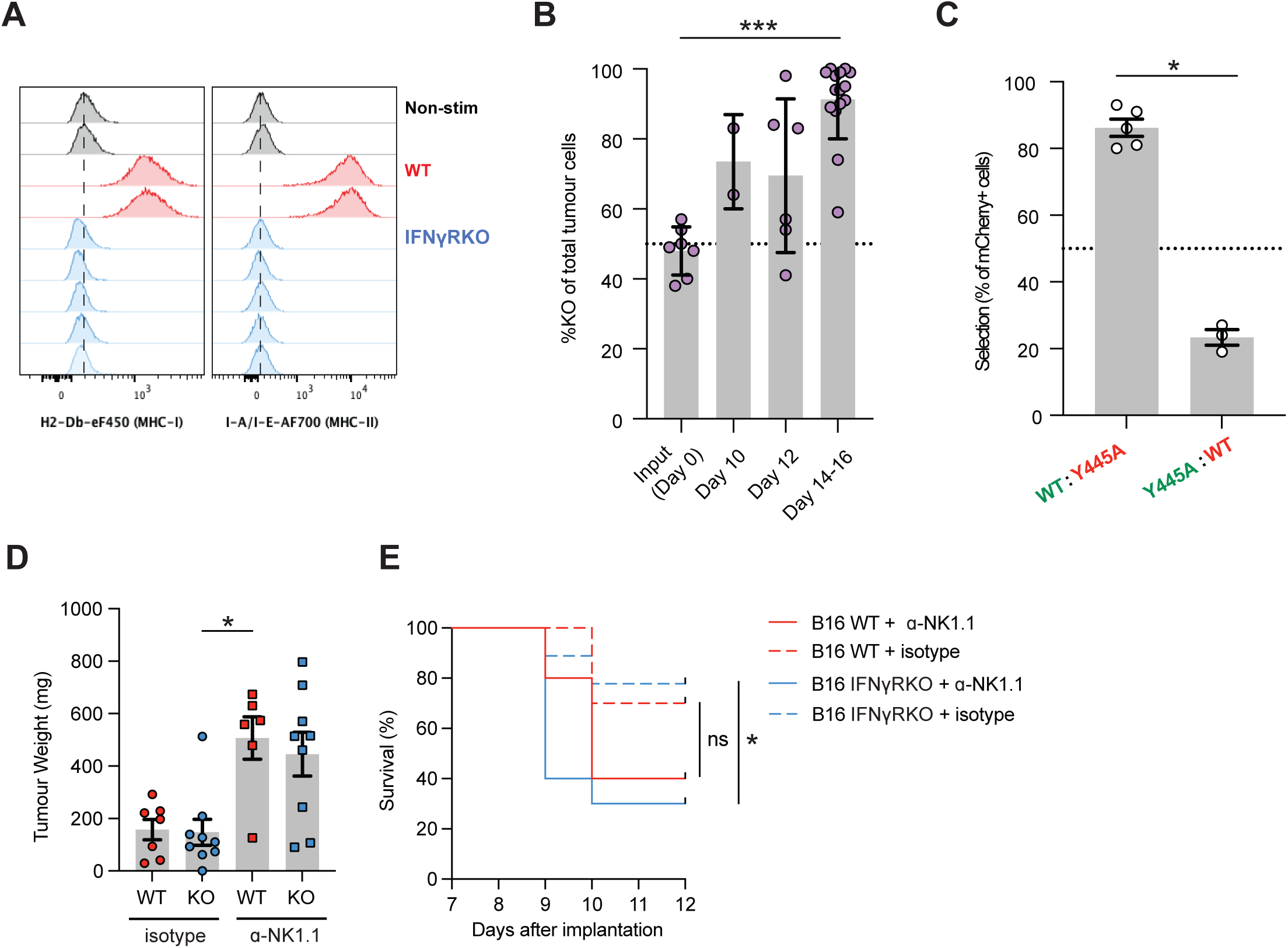
Validation of the B16-OVA IFNγRKO tumour models. (**A**) *In vitro* B16-OVA cells stimulated with 10ng/mL recombinant murine IFNγ for 48h, followed by analysis of MHC-I (H2-D^b^) and MHC-II I-A/I-E expression by flow cytometry. WT cells are shown in red, in duplicate samples, and five individual IFNγRKO clones are shown in blue. (**B**) Percentage of cells from WT/IFNγRKO admixed tumours at indicated time points following tumour engraftment, analysed by flow cytometry and gated on live CD45^-^ population. (**C**) Percentage of cells from tumours composed of B16-OVA WT and IFNγR Y445A mutant cells admixed 1:1 prior to engraftment. Green and red writing indicates that the indicated tumour cells express ZsGreen and mCherry, respectively. Tumours were analysed at day 14 post-engraftment. Data is from one experiment and n=8 total. (**D-E**) WT and IFNγRKO tumours were engrafted in WT mice treated with NK depleting antibodies prior to and following tumour engraftment. (**D**) Tumour volumes at day 12 post-engraftment. (**E**) Survival over time. Data show mean ± SEM with *p*-values by non-parametric Mann-Whitney *t* tests for comparisons between two groups, or Kruskal-Wallis tests between three groups with multiple comparisons correction using Dunn’s method. *p*≤0.05; \*\*\**p*≤0.001.

**Figure S3.**
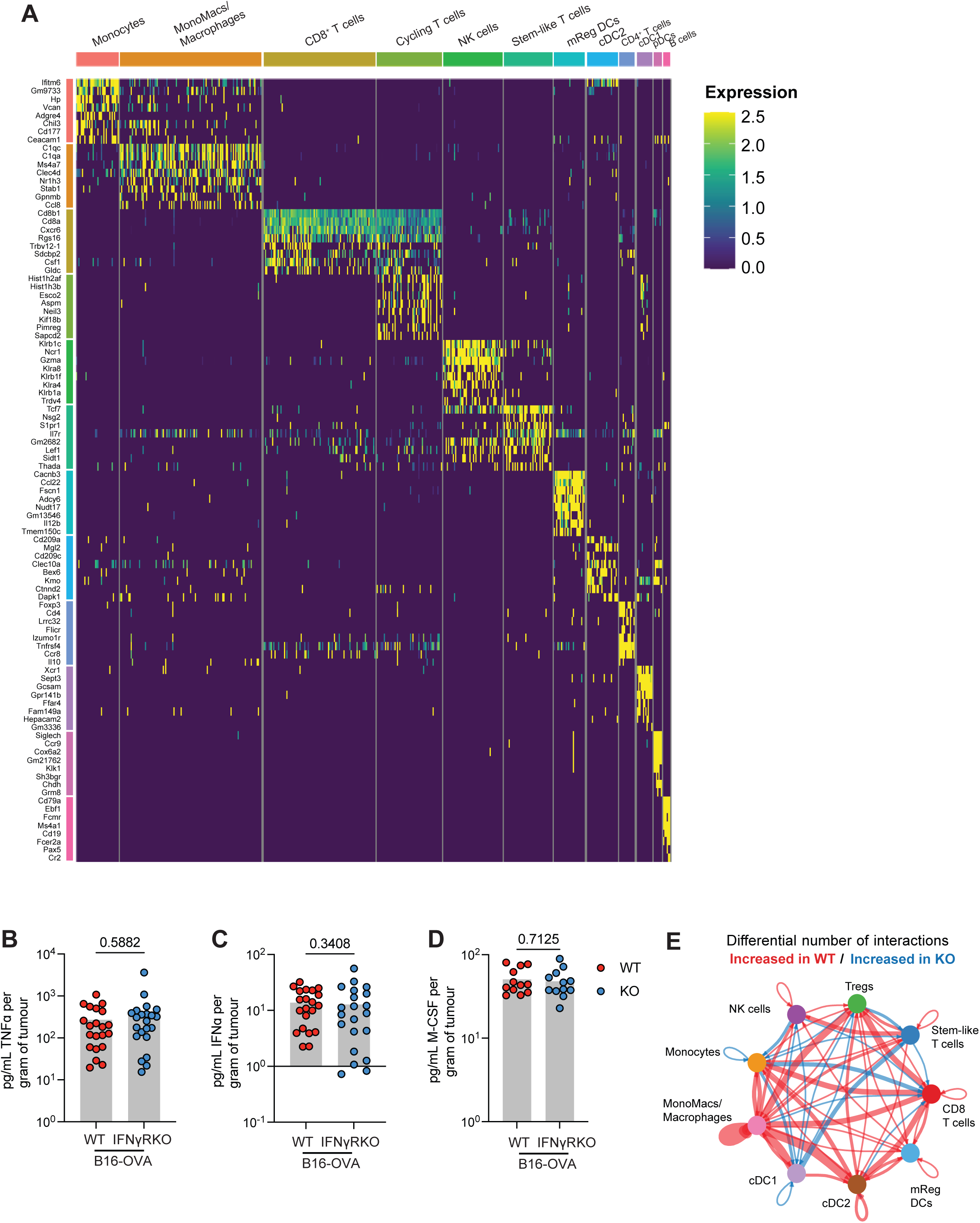
Characterisation of the B16-OVA IFNγRKO tumour landscape. (**A**) Heatmap of the top 8 genes expressed by unique clusters identified by scRNAseq from Fig.3. (**B-D**) Quantification of TNF (**B**), IFNɑ (**C**) and M-CSF (**D**) from tumour supernatant samples by Legendplex multiplex cytokine bead array analysis. Data show mean ± SEM with *p*-values by non-parametric Mann-Whitney *t* tests for comparisons between two groups. (**E**) Circle plot visualizing number of signalling interactions between immune populations from WT and IFNγRKO tumours. Vertices represent independent populations, and arrows indicate direction of signals sent, where broader lines represent increased quantity of signalling interactions.

**Figure S4.**
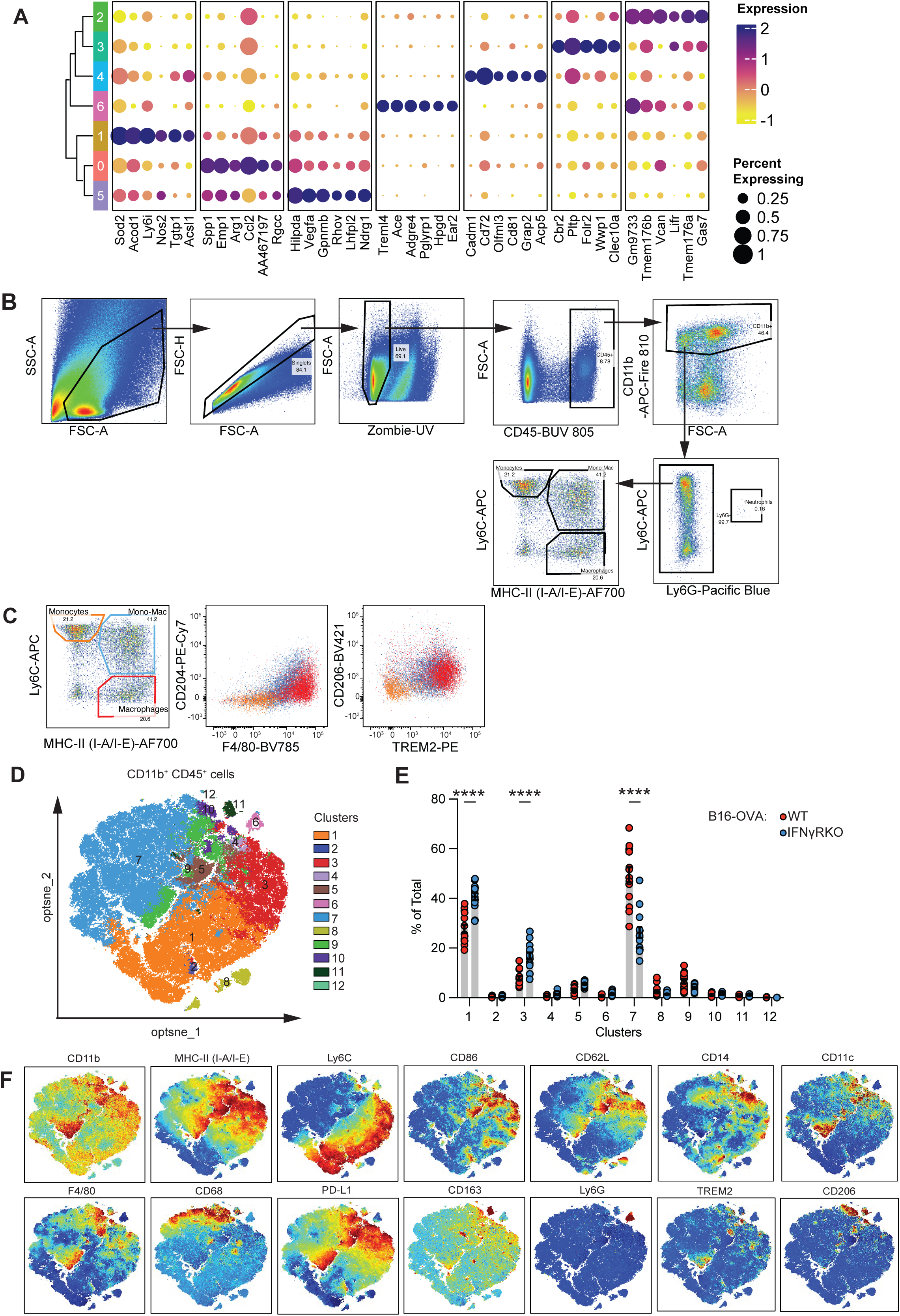
Adaptation of Tumour-infiltrating myeloid populations in B16-OVA IFNγRKO tumours. (**A**) Clustered dot plot of myeloid subpopulations from scRNAseq with dots coloured by top 6 genes for each cluster, and size of dots representing the percent of cells expression that gene. (**B-F**) WT or IFNγRKO tumours were engrafted in WT mice and analysed by flow cytometry at endpoint. (**B**) Gating example for analysis of monocyte and macrophage populations in the tumour. (**C**) Representative dot plot for Ly6C and MHC-II expression which is gated by monocytes (Ly6C^+^/MHC-II^-^), mono-macs (Ly6C^+^/MHC-II^+^) and macrophages (Ly6C^-^/MHC-II^+^), with expression of F4/80, CD204, CD206 and TREM2 shown for each subpopulation by gate colour. (**D**) optSNE plots of spectral flow cytometry data which shows clusters identified by OMIQ software analysis. (**E**) Relative quantification of clusters 1-12 of WT and IFNγRKO tumours. (**F**) Pseudocolour plots of individual marker expression, mapped to the optSNE projections. Colours shown (high expression in red, low expression in blue) are respective to each marker. Plots in D-F are from n=8-10 tumours for WT and IFNγRKO from one experiment, and representative of three independent experiments.

**Figure S5.**
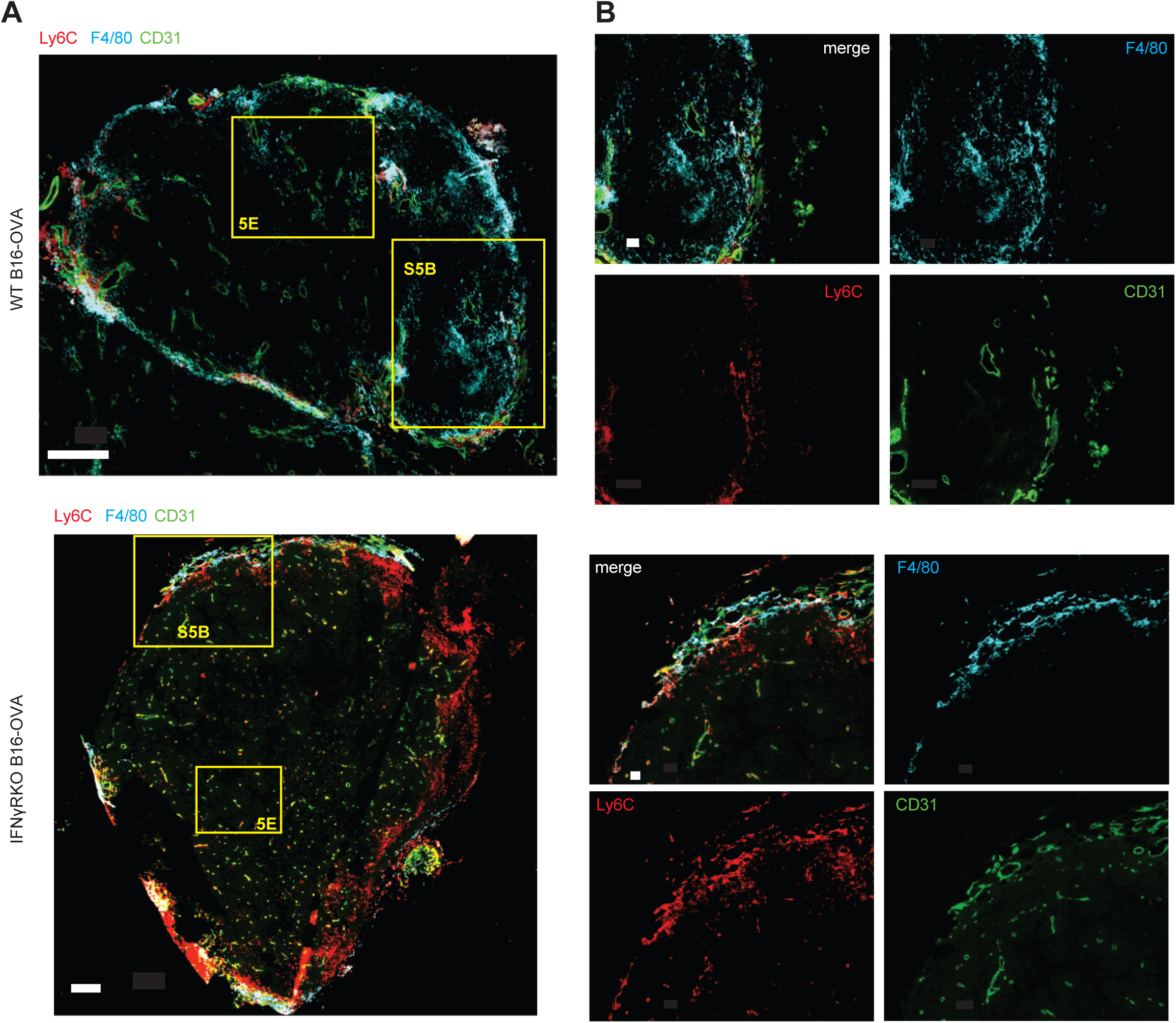
Intra-tumoural localisation of myeloid cells. WT or IFNγRKO tumours were engrafted in WT mice and imaged between day 11 to 13. Localisation of Ly6C (red), F4/80 (Cyan) relative to blood vessels (CD31, green) in whole tumour (**A**) or at the margin (**B**). Scale bar = 300um (**A**) and 50um (**B**). Yellow squares in (**A**) represent the location from zooms in (B) and Fig.5E.

**Figure S6.**
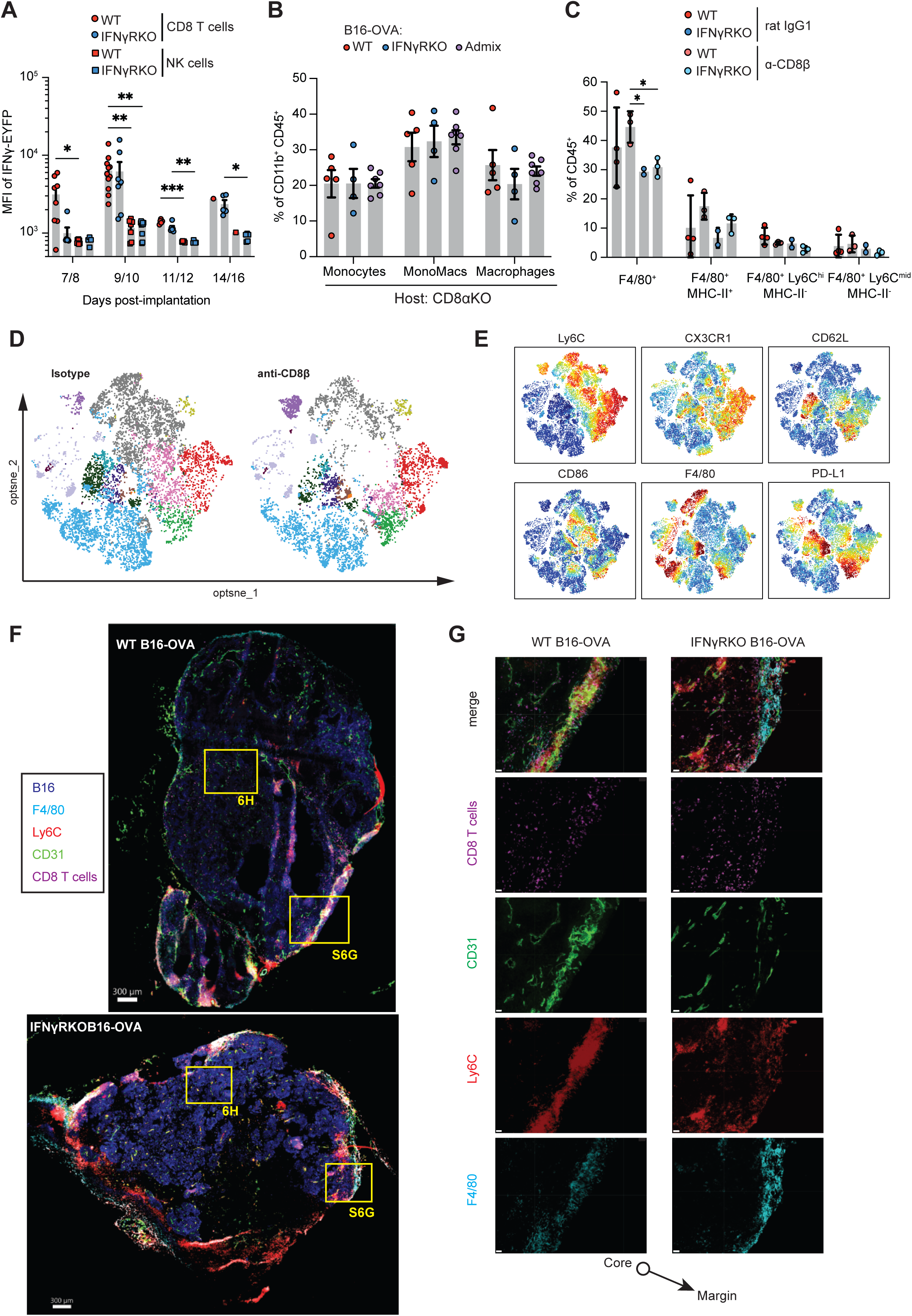
Role of CD8+ T cells for controlling IFNγRKO tumours. (**A**) WT or IFNγRKO tumours were engrafted in GREAT mice and harvested when indicated. IFNγ expression was quantified EYFP gMFI in tumour-infiltrating CD8^+^ T cells and NK cells. Data are pooled from four independent experiments, with timepoints varying between experiments. (**B**) Infiltration of myeloid populations relative to total CD11b^+^CD45^+^ cells in WT, IFNγRKO or admixed tumours engrafted into CD8aKO mice. (**C-E**) Antibody depletion of CD8^+^ T cells using anti-CD8b prior to and following tumour engraftment. (**C**) Frequency of specific macrophage (F4/80^+^) subsets following CD8^+^ T cell depletion was analysed by flow cytometry 13 days post-engraftment. (**D**) optSNE plots of high-dimension flow cytometry data of CD11b^+^ CD45^+^ tumour infiltrating cells from anti-CD8b and isotype-treated tumour-bearing mice. (**E**) Pseudocolour overlays of marker expression by different cell clusters. optSNE plots are concatenated data from three biological replicates for each condition. Data is from one experiment. (**F-G**) WT or IFNγRKO tumours were engrafted in WT mice and imaged between Day 11 to 13. Localisation of Ly6C (red), F4/80 (cyan), and CD8^+^ T cells (magenta) relative to blood vessels (CD31, green) in whole tumour (**F**) or at the margin (**G**). Scale bar = 300um (**F**) and 30um (**G**). Yellow squares in (**F**) represent the location from zooms in (**B**) and Fig.6G.

**Figure S7.**
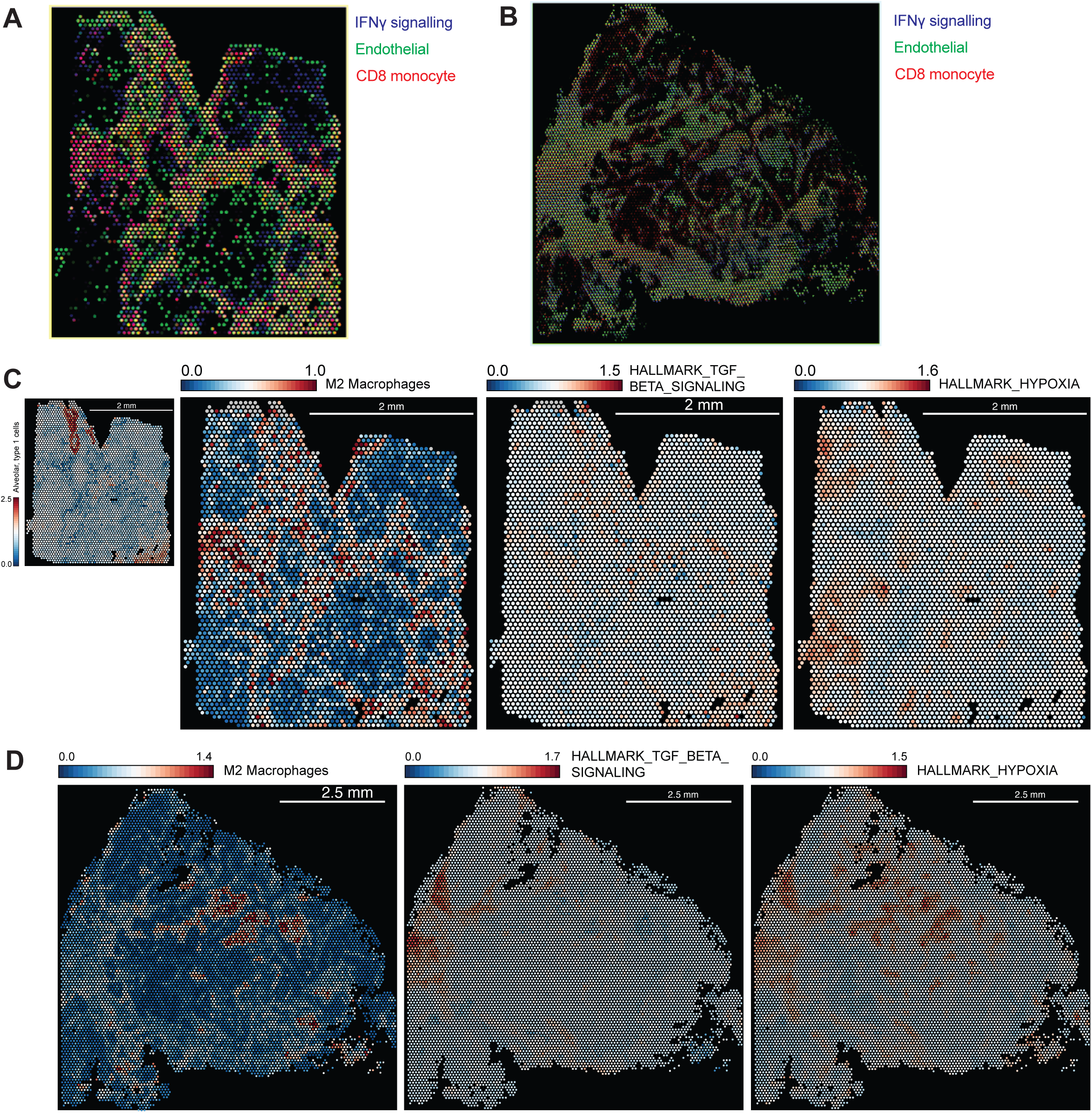
Gene signature analysis of 10X Genomics Visium spatial datasets. (**A-B**) Overlay of hallmark IFNγ response (blue), CD8-monocyte (red), and endothelial cell (green) gene signatures from 10X Genomics Visium datasets of human lung squamous cell carcinoma (**B**) or colon adenocarcinoma (**C**) samples from Fig.7A-B. (**C-D**) Analysis of 10X Genomics Visium datasets for M2 macrophages, hallmark TGFbeta signalling and hypoxia gene signatures in human lung squamous cell carcinoma (**C**) or colon adenocarcinoma (**D**) samples. Gene set expression is indicated by heatmap, where colours represent log-normalized average expression.

**Supplementary Dataset 1.** Gene signatures used in the study and sources.

## Notes

### Competing Interest Statement

The authors have declared no competing interest.

